# Real-Time Ratiometric pH Imaging of Macrophage Lysosomes Using the Novel pH-sensitive Probe ApHID

**DOI:** 10.1101/2024.01.20.576118

**Authors:** Santiago Solé-Domènech, Pradeep Kumar Singh, J. David Warren, Frederick R. Maxfield

## Abstract

Lysosomes actively regulate their lumenal pH, which is necessary for optimal enzymatic activity. Endocytic processes are involved in many diseases, including Alzheimer’s disease, in which sub-optimal lysosomal function has been reported. To measure acidification, pH-sensitive probes can be delivered to endosomes and lysosomes using labeled dextran polymers or proteins. However, many commercially available probes have limited sensitivity in the acidic range of lysosomes, and their fluorescence is subject to enzymatic degradation and photobleaching. Herein, we describe the preparation, characterization, and use of a novel pH-sensitive probe, *ApHID*, a green-emitting fluorescent dye with optimal dynamic range within the acidity of endosomes and lysosomes. ApHID has a pKa near 5, increasing brightness with acidity, and it is robustly resistant to oxidation and photobleaching. We used ApHID ratiometric imaging to measure lysosomal pH in macrophages, yielding virtually identical results when compared with fluorescein and Oregon Green. Overall, *ApHID* circumvents limitations presented by most commercially available pH-sensitive probes and can be useful in demanding imaging applications such as intravital imaging of tissues.

## MOTIVATION

Commercially available fluorescence probes have a limited capacity to sense LE/Ly pH. Fluorescein, which has been widely used to measure endosomal pH, has a pKa of 6.5, and its brightness decreases with acidity, which limits its sensitivity in the acidic range of LE/Lys (pH 4.5-5.5). Also, fluorescein undergoes rapid photobleaching, which limits its use in applications requiring extended imaging. In the past two decades, several pH-sensitive probes were developed commercially, including Oregon Green, pHrodo red, pHrodo deep red, and pHrodo green, among others, with improved fluorescence dynamic range in the acidic spectrum of endosomes. Oregon Green has a pKa of 4.7, and its dynamic range between pH 4 and 6 is better than that of fluorescein. However, its brightness also decreases with acidity, limiting its sensitivity. pHrodo red and pHrodo green brightness increases with acidity, but the probes have a pKa of 6.5, which, like fluorescein, limits their sensitivity in the acidity range of LE/Lys. The pHrodo deep red probe, which was developed recently, has a pKa of 5 and a good dynamic range in the pH reach of 4.0-5.5, but its fluorescence needs to be detected in the far-red spectrum, and its brightness is modest, which limits its uses. For prolonged fluorescence excitation or intravital imaging of tissues, which can cause extensive photobleaching and light scattering, bright probes with robust fluorescence, resistance to chemical modification, and excellent fluorescence dynamic range between pH 4.0 and 6.0 are required.

With that in mind, we designed a novel, BODIPY-based pH-sensitive probe, called *ApHID*, a green-emitting dye with a pKa of ∼5 that brightens up with increasing acidity. ApHID shows a remarkable dynamic range in the pH range of 4.0 to 6.0 and is robustly resistant to chemical modifications and photobleaching. The probe can be attached to dextran polymers, together with a second, pH-independent probe, for labeling of acidic compartments and ratiometric imaging.

## INTRODUCTION

Mammalian cells use a variety of endocytic mechanisms to internalize small molecules, macromolecules, and particles that are delivered to specific sealed organelles ^1^. Late endosomes and lysosomes (LE/Ly) are membrane-bound vesicles containing more than 60 different hydrolases and more than 100 membrane proteins, which constitute the degradative organelles of the endocytic system ^2^. These organelles have the capacity to tightly regulate their intraluminal pH, which is required for maintaining optimal enzymatic activity ^3^. In most cells, during the 30-60 minutes following internalization, ligands encounter an increasingly acidic environment ranging from about pH 6 in sorting endosomes to a pH of 4.5-5 in lysosomes ^4^. The main regulator of vesicular pH is the V-ATPase complex, with its different subunits playing a role in the regulation of membrane potential and vesicular pH as well as organelle function ^5^.

Endocytic processes play a number of roles in many diseases, including Alzheimer’s disease (AD) and atherosclerosis, and lysosomal enzymatic deficiencies lead to lysosomal storage disorders such as Tay-Sachs disease and ceroid lipofuscinosis ^6^. LE/Ly membrane permeabilization can be caused by a variety of factors ^7^, and AD’s fibrillar amyloid-beta (Aβ) has been reported to damage LE/Lys and cause enzyme leakage ^8^. During AD pathogenesis, a deficiency in endolysosomal acidification has been shown to block autophagic flux, causing neurons to fill with undigested autophagic cargo leading to extensive cellular damage ^9^. Also, it has been hypothesized that aging might diminish overall endolysosomal function, which could exacerbate neurodegenerative conditions ^10^. Microglia, the immune cells of the brain, can become activated in parts of the AD brain, causing damage to neurons while probably inefficiently digesting fibrillar Aβ due to insufficient LE/Ly function ^11,12^. It is, therefore, important to find tools to visualize LE/Ly compartments and measure their acidification both in cell culture and *in vivo*. Novel methodology in this direction would aid in the understanding of pathogenic processes for which our knowledge is still limited.

LE/Ly pH can be quantified in cells by fluorescence microscopy imaging of organelles labeled with pH-sensitive dyes, which can be delivered to the organelles using labeled dextrans or proteins ^1^. Once incorporated into the endocytic system, dextrans or proteins labeled with pH-sensitive probes reach endocytic compartments, where they serve as pH sensors. Organelle pH can be determined with precision by ratiometric imaging and ratios measured can be interpolated to pH values using a ratio-to-pH calibration prepared using fixed cells. Sorting endosomes have a luminal pH of 5.9-6.0, whereas LE/Lys have a pH of 4.5-5.5 ^1,13^. Hence, to accurately measure pH in these organelles, a probe must be most sensitive to the pH range of 4.0-6.0. In 1978, Okhuma and Poole measured LE/Ly pH in macrophages using fluorescein-dextrans ^14^. We and others have used ratiometric pH imaging of dextrans or proteins labeled with various fluorescence probes to measure lysosomal and pH in many types of cells ^15–25^. We also measured the acidity of extracellular degradative compartments -*lysosomal synapses-*, formed by macrophages and microglia interacting with large aggregates of LDL or extracellular Aβ deposits during digestive exophagy, respectively, using ratiometric pH imaging ^26,27^.

In the present work, we describe the preparation, characterization, and use of a novel pH-sensitive probe, which we have called Acid pH Indicator Dye, *ApHID.* With a pKa near 5 and an emission maximum at 515 nm, ApHID displays excellent dynamic range in the pH range of acidic organelles, and its brightness increases with acidity. ApHID fluorescence is highly resistant to photobleaching, and once incorporated into LE/Ly compartments, its fluorescence is stable and resistant to oxidation. We used ratiometric pH imaging to measure LE/Ly pH comparatively in macrophages using dextrans labeled with ApHID, fluorescein or Oregon Green and the pH-independent dye Alexa 647 and obtained virtually identical pH readings for all probes. Overall, we believe that ApHID constitutes a promising tool that circumvents some limitations presented by commercially available pH probes.

## RESULTS

### Design of a pH-sensitive probe to measure acidity in endosomes and lysosomes

*ApHID* is composed of a BODIPY core with two flanking amide substitutions and an electrophilic nature (Fig. 1A). The BODIPY core is attached to an aniline moiety acting as an electron donor. The electron donor propensity of the aniline moiety determines the range of pH sensitivity of the molecule, and it can be conveniently modulated by attaching different alkyl groups to the nitrogen of the *N,N*-dialkyl-*o*-toluidine moiety. Based on a previous study by Maeda and collaborators^28^, we attached two methyl groups to the nitrogen of *N*,*N*-dialkyl-*o*-toluidine moieties (Fig. 1A). To increase water solubility of the otherwise insoluble structure, we attached a 10-carbon polyethylene glycol chain (PEG4) to one of the amide groups. The distal end of the PEG4 chain was derivatized with an *N*-hydroxysuccinimidyl ester group, allowing for the labeling of proteins and other molecules such as amino-dextrans via reaction with primary amines (group R in Fig. 1A).

**Figure 1.**
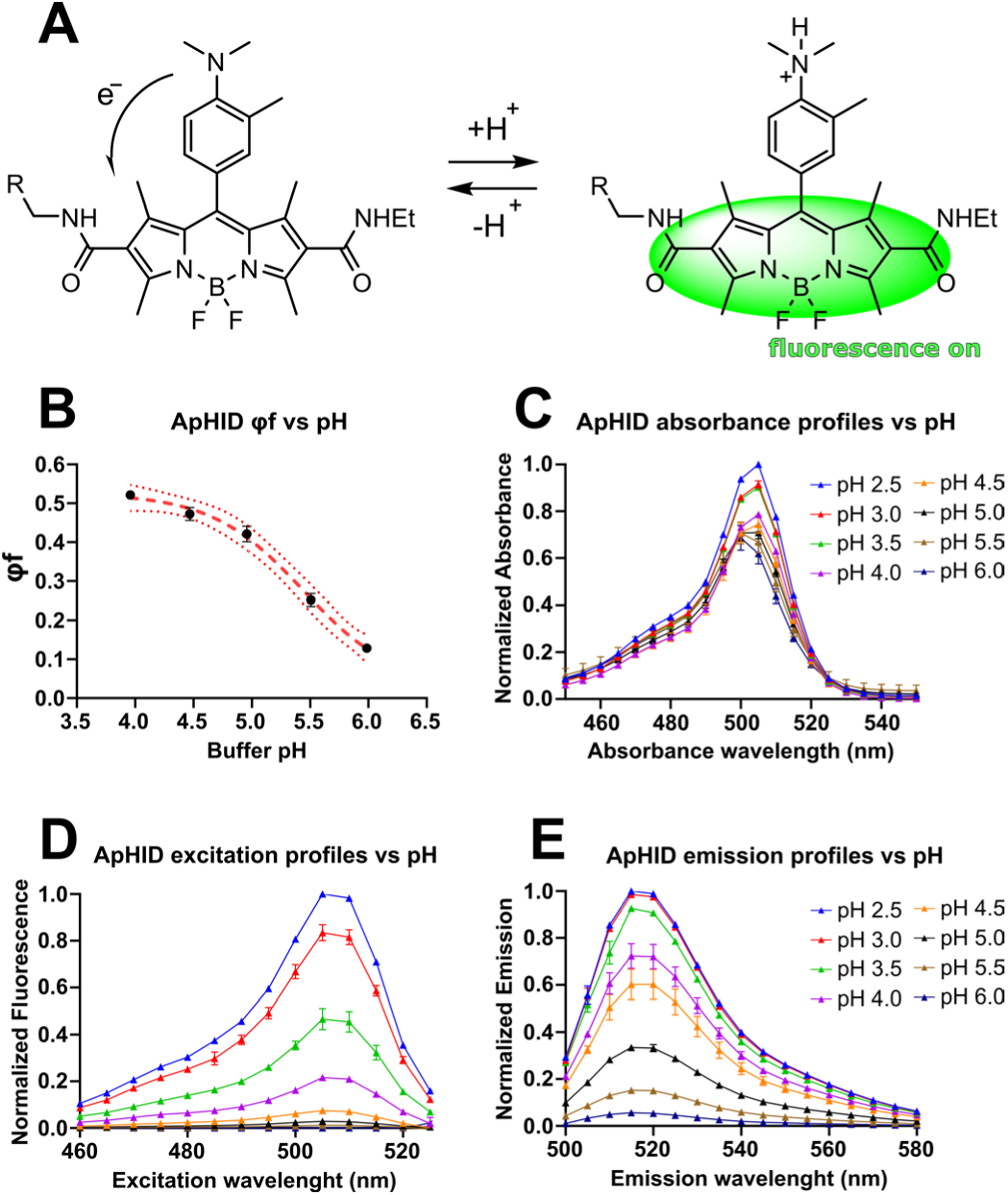
On-off switch and spectroscopic properties of ApHID, **(A)** ApHID structure and on-off fluorescence switch. ApHID contains an aniline moiety acting as an electron donor at alkaline pH, which quenches fluorescence. The group ‘R’ consists of a 10-carbon polyethylene glycol chain (PEG4), which ends in a succinimidyl ester group that confers increased solubility in water and allows for derivatization of molecules containing primary amines such as proteins or amino-dextran polymers. **(B)** Quantum yield of the hydrolyzed NHS ester form of ApHID measured in various pH-adjusted buffers, using fluorescein in 0.01 M NaOH (pH 12) as a standard. Quantum yield was plotted against pH, yielding a titration that was fit to a 4-component sigmoid with -log IC_50_ (pKa) of 5.4. The maximal extinction coefficient and quantum yield of ApHID, measured in a subsequent experiment in pH 3.0 buffer, are 0.64 and 99,710 M^-1^ cm^-1^, respectively. **(C-E)** Absorbance (C), excitation (D), and emission (E) spectra plotted against buffer pH, measured for a 0.04 mg/mL dilution of 10 KDa amino-dextran labeled with ApHID at a 1.6:1 molar ratio, in pH-adjusted buffers. Two independent measurements were completed for each experiment. Geometrical objects and bars indicate averages ±SEM.

The quantum yield (QY) for ApHID is 0.64 [measured relative to 5/6-carboxyfluorescein carboxylic acid (fluorescein) as a standard in 0.1 M NaOH], and its extinction coefficient is 99,700 M^-1^ cm^-1^, both measured in citrate buffer at pH 3.0. The pKa value of ApHID (hydrolyzed NHS ester form) in solution is 5.4, based on its pH-dependent QY profile (Fig. 1B). The UV-visible absorption spectra of ApHID remain similar between pH 4.0 to pH 6.0 (Fig. 1C), but fluorescence emission spectra increase sharply in amplitude with increasing acidity (Fig. 1D). Excitation spectra were also pH-dependent, with an excitation maximum at 506 nm (Fig 1E).

### ApHID fluorescence and pKa remain stable in the presence of •OH radicals, protein, or salts in solution

We compared the spectroscopic properties of ApHID with those of commercially available pH-sensitive probes, namely LysoSensor yellow/blue, fluorescein, and Oregon Green. Since LysoSensor is commercially available attached to 10 KDa dextran polymers, and since we will be using derivatized dextrans in subsequent experiments, we carried out our measurements using probes attached to dextran polymers. Dextran labeling can be achieved by reacting the fluorophores, attached to an *N*-hydroxysuccinimidyl ester (NHS) group, with polymers previously derivatized with primary amines (amino-dextrans).

Fluorophores attached to 10 KDa dextrans were solubilized in buffers with pH adjusted between 1.5 and 8.5 (see Methods), and fluorescence and absorbance were measured and plotted against buffer pH. The resulting absorbance profiles for LysoSensor, Oregon Green and fluorescein could be fit to a sigmoidal curve, but ApHID absorbance increased monotonically with increasing acidity (Fig. 2A). The fluorescence emission profiles for all probes, including ApHID, fit a sigmoidal curve (Fig. 2B). ApHID fluorescence at pH 4.0 is almost 12 times greater relative to pH 6.0, while LysoSensor is 7 times brighter. Fluorescein and Oregon Green fluorescence decreases with acidity, being 3 and 6 times brighter at pH 6.0 relative to pH 4.0, respectively (Fig. 2B). By this analysis, ApHID shows the greatest dynamic range at the pH range of LE/Lys.

**Figure 2.**
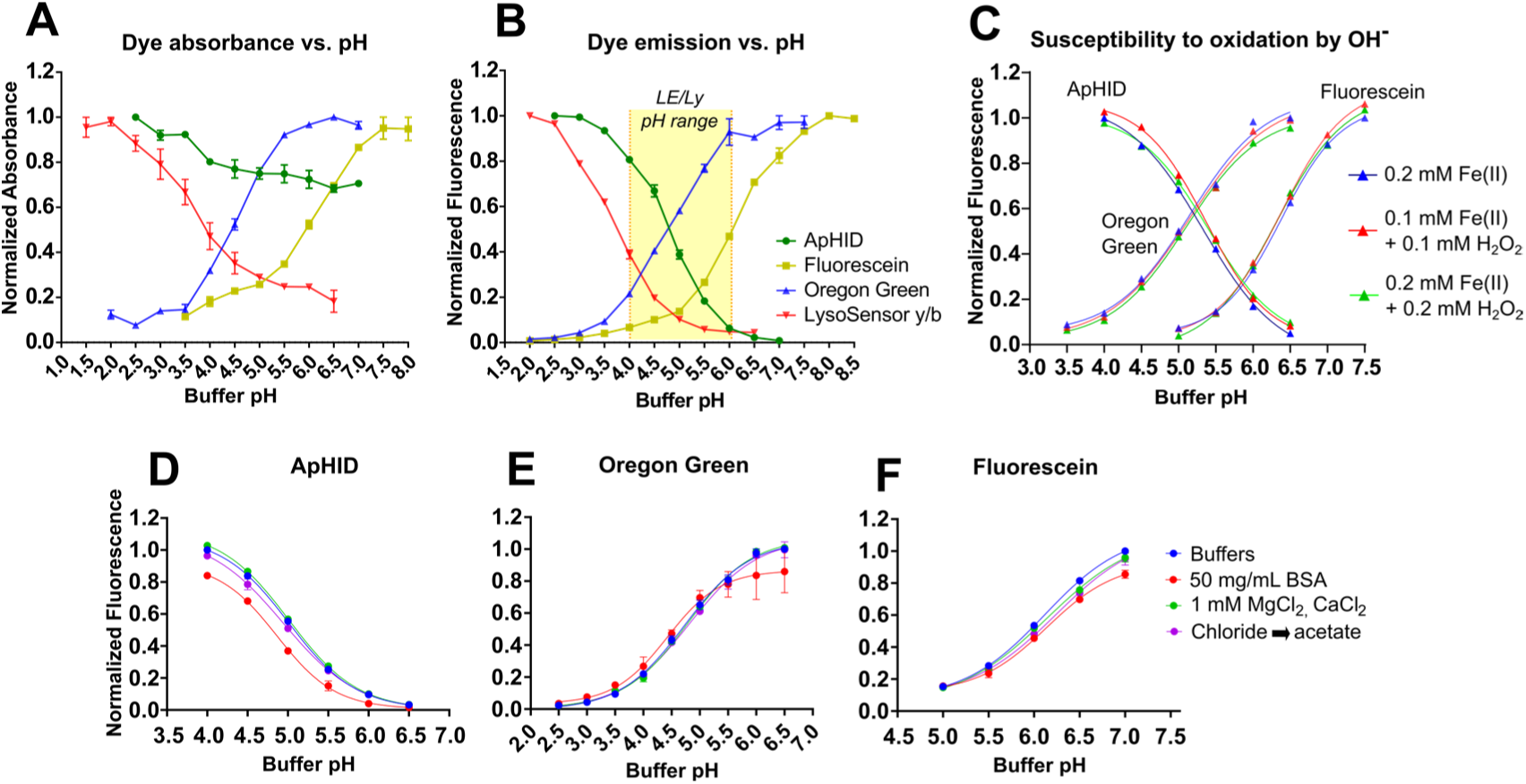
ApHID fluorescence and pKa are not affected by oxidation caused by •OH radicals, protein, or various salt concentrations in solution. **(A, B)** ApHID, fluorescein, Oregon Green, and LysoSensor y/b absorbance (A) and emission (B) profiles vs buffer pH, measured in solution. Each dye was attached to 10 KDa amino-dextran polymers and diluted to 0.02 mg/mL in various pH-adjusted buffers. ApHID absorbance decreases monotonically with increasing pH, and emission increases strongly with increasing acidity. **(C)** ApHID, Oregon Green, and fluorescein (hydrolyzed NHS ester forms) fluorescence titrations against buffer pH in the presence of various amounts of hydroxyl radical (•OH) generated by mixing various amounts of ferrous perchlorate (II) and H_2_O_2_ in solution, incubated at 37 °C for 20h. **(D-F)** ApHID (D), Oregon Green (E), and fluorescein (F) dyes attached to 10 KDa amino-dextrans were diluted to 0.04 mg/mL in pH-adjusted buffers enriched either with 50 mg/mL BSA, 1 mM CaCl_2_, and 1 mM MgCl_2_ or sodium acetate instead of sodium chloride and incubated at 37 °C for 20h. For all experiments, measurements were repeated twice. Normalized fluorescence intensities were plotted against buffer pH resulting in various titrations and fit to 4-component sigmoidal curves. In all panels, geometrical objects and bars indicate average ±SEM.

Dextrans derivatized with fluorophores can be used to label and track LE/Lys in different cell types. Depending on the cell type, the probes may be subjected to reactive oxygen species generated intracellularly ^29^, which could cause chemical modifications that affect spectroscopic properties. The most reactive ROS species is the hydroxyl radical (•OH) ^30^. To test the effects of •OH on ApHID, fluorescein, and Oregon Green fluorescence, hydrolyzed NHS ester forms of the probes were added to PBS solutions containing 100 or 200 µM •OH and incubated for 24h at 37 °C. The •OH radicals were generated by Fenton Reaction by mixing ferrous perchlorate, Fe(II), with H2O2 in solution. As a control, we incubated the probes with ferrous perchlorate in the absence of H2O2. The intracellular concentration of H2O2 in various cell types, under physiological conditions, is within the range of 1-10 uM ^31,32^. However, ROS concentrations in some cancer microenvironments can be as high as 100 µM, depending on the activation status and metabolic state of the cells ^33^. We conducted our assay with conditions that would ensure •OH concentrations above these reported levels. Following reaction, the mixtures were diluted in buffers with pH ranging between 4 and 7.5, and fluorescence was measured and plotted against buffer pH (Fig. 2C). Only small variations in pKa were observed, indicating that the probes were resistant to oxidation by •OH for at least 24h (Table 1).

**Table 1.** Log IC_50_ (pKa) values ± SEM calculated from sigmoidal curves fit to the titrations shown in Figs. 2C-2F. Buffers with pH adjusted between 1.5 and 8.5 (see Methods) alone or enriched in either BSA, CaCl_2_ + MgCl_2_, or sodium acetate (as a substitutive for sodium chloride) were tested on fluorescent probes attached to dextrans. Oxidation by •OH was tested on the unattached, hydrolyzed succinimidyl ester form of the dyes. Note that the pKa of the probes is generally lower when attached to dextrans. This has been reported previously for fluorescein ^36^. The experiments were repeated twice.

|  | ApHID | Oregon Green | Fluorescein |
| --- | --- | --- | --- |
| **pH-adjusted buffers** | 5.02 ± 0.02 | 4.74 ± 0.06 | 6.11 ± 0.01 |
| **Buffers + 50 mg/mL BSA** | 4.87 ± 0.01 | 4.41 ± 0.15 | 6.15 ± 0.04 |
| **Buffers + 1 mM MgCl<sub>2</sub>, CaCl<sub>2</sub>** | 5.02 ± 0.02 | 4.79 ± 0.02 | 6.17 ± 0.06 |
| **Buffers + Chloride → Acetate** | 4.94 ± 0.04 | 4.82 ± 0.03 | 6.26 ± 0.08 |
| **Buffers + 0.2 mM Fe(II)** | 5.32 ± 0.08 | 5.15 ± 0.01 | 6.38 ± 0.02 |
| **Buffers + 0.1 mM $\bullet$OH** | 5.36 ± 0.08 | 5.14 ± 0.02 | 6.38 ± 0.06 |
| **Buffers + 0.2 mM $\bullet$OH** | 5.39 ± 0.09 | 5.12 ± 0.06 | 6.34 ± 0.07 |

Fluorophores reaching acidic compartments might also be sensitive to the high protein ^2^ and salt ^34,35^ concentration present in the organelles. To test for this possibility, 10 KDa dextrans labeled with the probes were incubated in buffers containing 50 mg/mL BSA, 1 mM MgCl2 and CaCl2, or sodium acetate in the absence of sodium chloride for 24h at 37 °C, and fluorescence was quantified and plotted against buffer pH thereafter. The probes were generally stable in the presence of salts, but their fluorescence decreased in the presence of BSA (Fig. 2D-2F). The pKa of the probes did not register large variations over the various conditions tested (Table 1), but it was generally lower when attached to dextrans, relative to its unattached molecular form, which has been reported in the literature for fluorescein ^36^.

### ApHID is resistant to photobleaching

Live cell and intravital fluorescence imaging techniques conducted for extended periods of time benefit from probes that can withstand prolonged, intense photoexcitation while maintaining their fluorescence properties. We determined ApHID resistance to photobleaching, induced by a 488 nm laser, over time and compared it to that of fluorescein and Oregon Green. J774 macrophages were loaded with 70 KDa dextrans derivatized with either probe at 2:1 molar ratio followed by a 4h chase to ensure localization in LE/LY. Cells were then fixed in PFA and incubated in 50 mM TRIS maleate pH 5.0 buffer (for ApHID and Oregon Green) or 1X PBS pH 7.4 buffer (for fluorescein) containing methylamine hydrochloride, sodium acetate, nigericin and monensin for 5 mins at 37 °C to ensure buffer equilibration across membranes. Once equilibrated, cells were imaged in a confocal microscope while irradiated with a 488 nm laser for 0.5 seconds per cycle (50 cycles in total) with 1 second intervals between irradiation pulses, and images were acquired after each cycle. The laser was adjusted to yield 5.2 µW power at the front element of the 40X objective. Fluorescence intensity for each dye and cycle (F) was normalized to fluorescence on cycle 1 (F0) and plotted as F/F0 against irradiation time (Fig. 3G). At the end of the photobleaching experiment, ApHID fluorescence intensity in LE/Lys (Fig. 3B, 3E) had decreased by 12%, whereas that of fluorescein (Fig. 3B, 3E) and Oregon Green (3C, 3F) had decreased by 83% and 82%, respectively (Fig. 3G). These results indicate that ApHID is highly resistant to photobleaching. See Suppl. Table ST1 for statistics.

**Figure 3.**
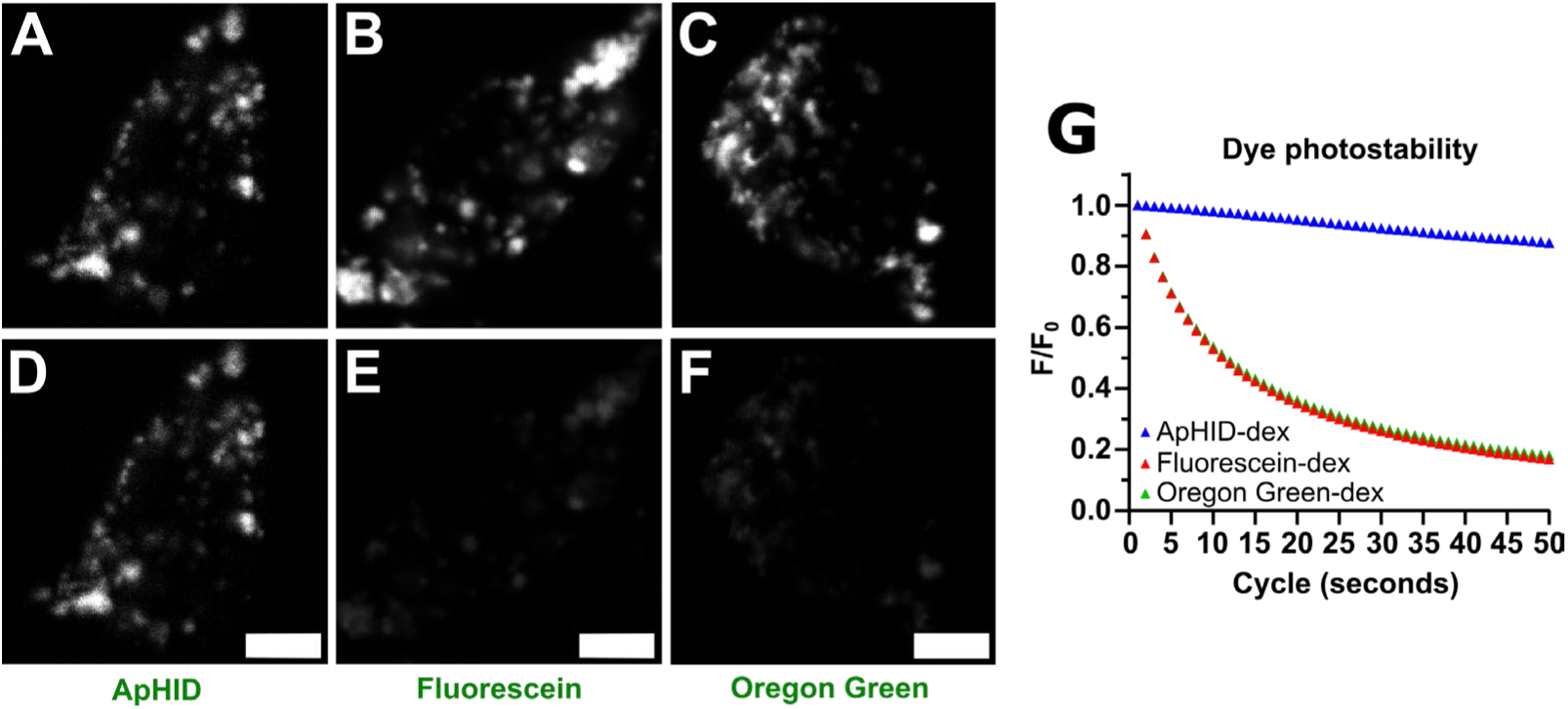
ApHID is highly resistant to laser-induced photobleaching. **(A-F)** J774 macrophages, imaged by confocal microscopy using a 40X objective, with LE/Lys labeled with 70 KDa dextran polymers derivatized with ApHID (A, D), fluorescein (B, E) or Oregon Green (C, F) at approx. 2:1 molar ratio, prior to (A, B, C) or after (D, E, F) exposure to laser irradiation using a 35 mW, 488 nm laser line for 50 half-second pulses. Cells were fixed with 0.5% PFA prior to imaging. ApHID and Oregon Green labeled cells were imaged in 50 mM TRIS maleate buffer adjusted to pH 5.0, and fluorescein was imaged in 1X PBS buffer adjusted to pH 7.4. Laser output was set to 30% (usual outputs during confocal imaging range between 0.5% and 4%), yielding 5.2 µW of power at the front element of the 40X objective used for imaging. Single confocal planes were irradiated for 50 cycles, with 1 second intervals between each cycle. Pixel dwell time was 33 µs. **(G)** Normalized fluorescence intensity was plotted against irradiation cycle for ApHID, fluorescein, and Oregon Green dextrans. The experiment was repeated twice; two dishes were measured per condition and 4 fields were imaged per dish for a total of 16 fields. Triangles indicate average fluorescence intensity for each irradiation cycle. The SD error bars fit within the symbols. See Suppl. Table ST1 for statistics. Scale bar: 5 µm.

### ApHID fluorescence and pKa remain stable with different degrees of amino-dextran derivatization and net charge

Amino-dextrans can be derivatized with a variety of pH-sensitive and pH-independent dyes (Fig. 4A, schematic representation, green and blue spheres respectively). Increasing dextran derivatization should increase overall brightness, which would be beneficial for applications with taxing light scattering or when extended imaging times and low laser powers are required. To test for the effect of dextran derivatization on ApHID fluorescence and pKa, we labeled 70 KDa polymers with different amounts of ApHID and a constant amount of Alexa 405 or Cy5•3xSO3^−^ dyes (pH-independent) and measured fluorescence against buffer pH in solution. Increasing derivatization with ApHID increases ApHID/Alexa405 and ApHID/Cy5•3xSO3^−^ ratios proportionally to their molar ratios (Fig. 4B, 4D). However, when normalized to pH 5.0 ratio, the resulting sigmoidal curves are not significantly different from each other (Fig. 4C, 4E). This indicates that ApHID fluorescence and pKa remain stable at a range of probe concentrations on the dextran. We also tested the effect of dextran net charge on ApHID pKa. To do so, we derivatized dextrans with a constant amount of ApHID and various amounts of Alexa 405, which carries three negatively charged sulfate groups (Fig. 4F). Increasing dextran negative charge with Alexa 405 increased ApHID pKa, but only at high negative charge density (Fig 4G). See Suppl. Table ST2 for statistics. These results indicate that ApHID fluorescence and pKa remain stable in a wide range of dextran derivatizations.

**Figure 4.**
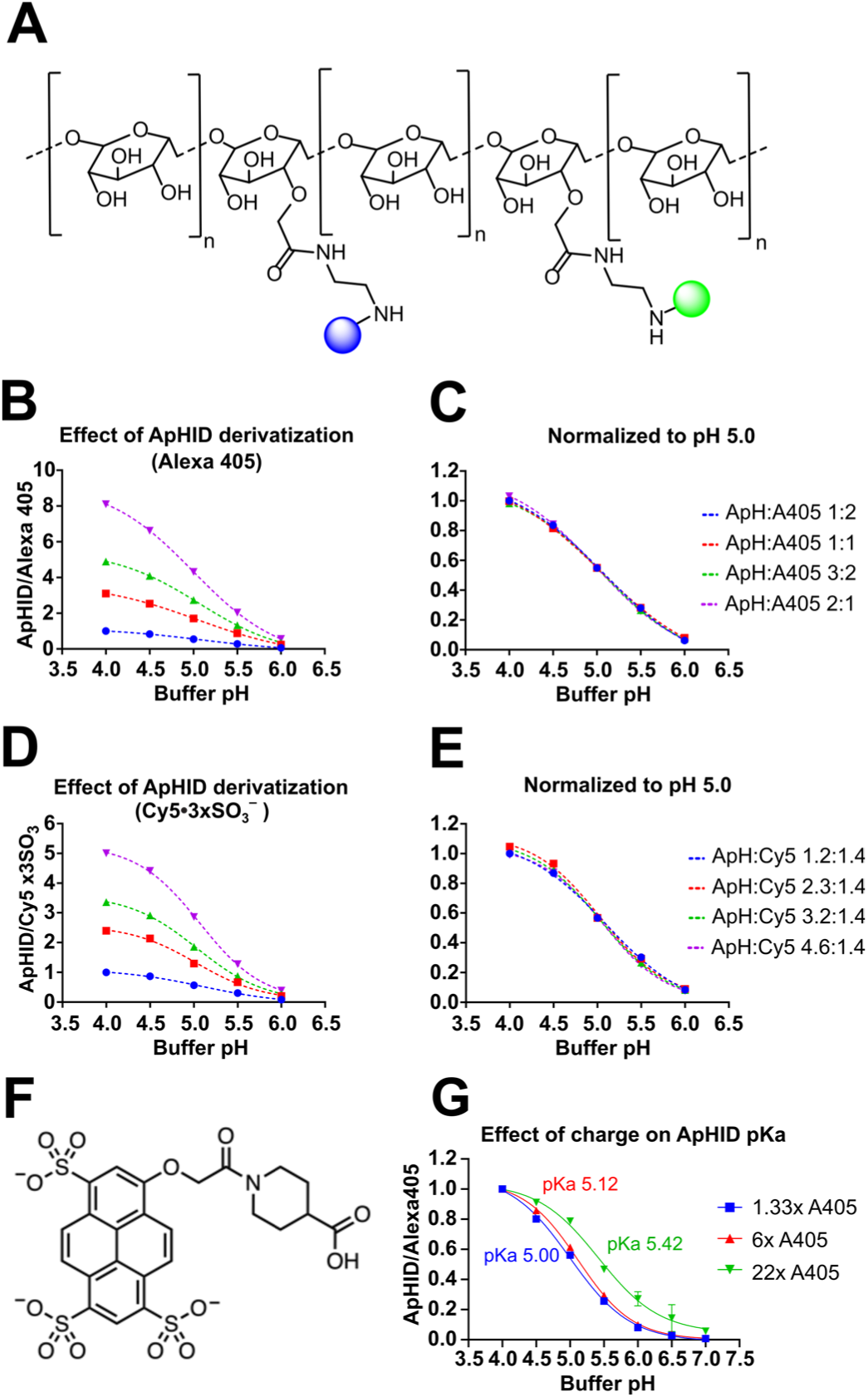
Effect of various amounts of other fluorophores and dextran polymer charge density on ApHID fluorescence and pKa profiles. **(A)** Representative structure of a dextran polymer containing lysine groups that can be derivatized with pH-sensitive (green spheres) and pH-independent (blue spheres) dyes. **(B-E)** ApHID/Alexa 405 (B, C) and ApHID/Cy5•3SO_3_^−^ (D, E) ratios measured for dextrans labeled with various amounts of ApHID, and a constant amount of either pH-independent dye, solubilized at 0.04 mg/mL in pH-adjusted buffers. Ratios were plotted against buffer pH. The titrations were either normalized to the ApHID/pH-independent ratio for the dextran containing the least amount of ApHID at pH 4.0 (B, D) or by the ApHID/pH-independent ratio for the dextran with the highest content in ApHID at pH 5.0 (C, E). **(F, G)** ApHID/Alexa405 ratios were measured for dextrans containing various amounts of Alexa 405 dye and a constant amount of ApHID. Each Alexa 405 molecule carries three negatively charged sulfate groups (F). Fluorescence ratios were plotted against buffer pH, and the various titrations were fit to a 4-component sigmoid from which -log IC_50_ (pKa) was calculated. pKa values are indicated next to each titration (G). See Suppl. Table ST2 for statistics. Experiments were repeated three times. Geometrical objects indicate averages. The SEM error bars fit within the symbols. Abbreviations: ‘A405’: Alexa Fluor 405; ‘Cy5’: Cy5•3SO_3_^−^; ‘ApH’: ApHID.

### Lysosomal pH measured in J774 macrophages using ApHID coincides with measurements done using fluorescein and Oregon Green and is stable over time

We have used ApHID previously to measure lysosomal pH in transfected HEK293 cells expressing TMEM106B mutants ^15^. We also used the probe to monitor the acidity of lysosomal synapses during digestive exophagy of model Aβ aggregates by primary microglial cells ^27^. In the current study, we use 70 KDa dextrans conjugated with ApHID and Alexa 647 (pH-independent) to measure lysosomal pH by ratiometric imaging. Many cell types endocytose dextrans and deliver them to their endosomal system.

We wanted to test whether ApHID, once incorporated in LE/Lys, reports lysosomal pH as reliably as fluorescein, a well-characterized probe that has been extensively used to measure pH in multiple cell types ^14,16^. Oregon Green was also tested for comparative purposes. Figure 5 shows a diagram with the described uses of ApHID and the current experimental approach to measure lysosomal pH using dextran polymers.

**Figure 5.**
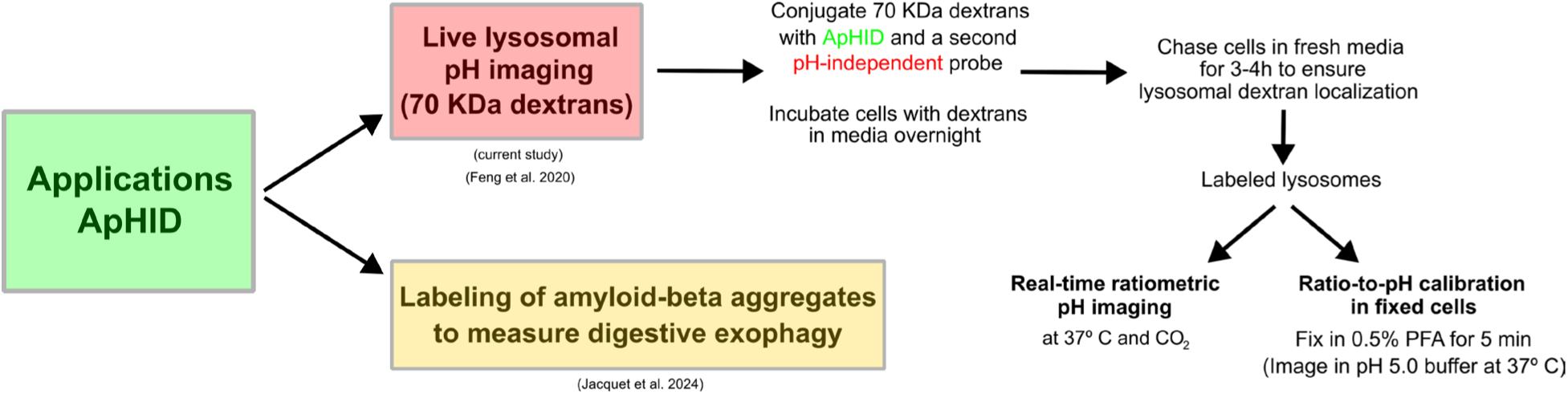
ApHID applications and summarized experimental approach to measure lysosomal pH using dextran polymers.

J774 macrophages were loaded overnight with 70 KDa dextrans labeled with either probe as well as Alexa 647 (pH-independent), followed by a 3h chase in fresh DMEM. After fixation, cells incubated in pH 4.5 buffer showed strong ApHID fluorescence, which decreased as the cells were equilibrated with increasing buffer alkalinity (Fig. 6A-D). To quantify pH-sensitive/Alexa 647 ratios in LE/Lys, an intensity threshold was applied to the pH-independent channel (Alexa 647). A mask was then generated and transferred to the pH-dependent channel. Integrated intensity was measured for each masked channel, and pH ratios were calculated for each LE/Ly or field imaged. ApHID/Alexa 647 ratios measured in fixed cells were plotted against buffer pH, and the resulting titration was fit to a 4-component sigmoidal curve and compared with a titration of the same dextran, measured in solution (Fig 6E). The sigmoidal curve corresponding to the titration in LE/Lys was practically identical to that in solution, indicating that ApHID and Alexa 647 are stable in LE/Lys for extended times.

**Figure 6.**
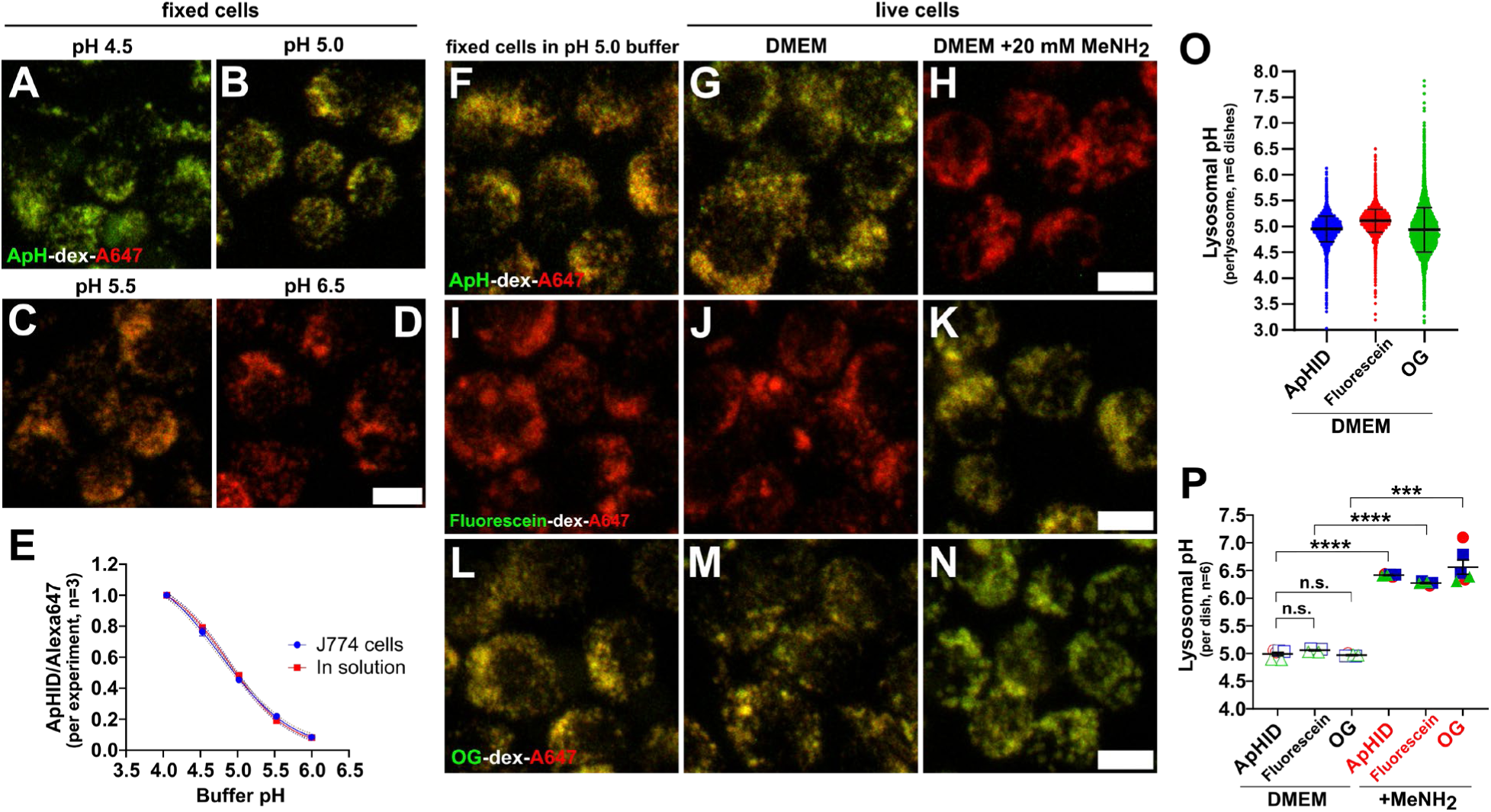
Lysosomal pH in J774 macrophages reported by ApHID matches that reported by fluorescein and Oregon Green. **(A-E)** Calibration confocal microscopy images of late endosomal and lysosomal (LE/Ly) compartments in J774 macrophages, loaded with fluorescent dextrans that were labeled with ApHID and Alexa 647. Cells were incubated with 0.5 mg/mL dextrans overnight and chased for 3h in fresh DMEM the following morning. Following chase, cells were fixed in 0.5% PFA and incubated for 20 min in pH 4.0 and pH 4.5 buffer or 30 min in pH 5.0-6.5 buffers at 27 °C, followed by imaging by confocal microscopy (A-D). ApHID/Alexa 647 ratios were calculated for each field and plotted against buffer pH. The resulting titration was fit to a 4-component sigmoidal curve (E, blue-dotted line) and compared with a titration of the same dextrans measured in solution using a spectrofluorometer, also at 27 °C (E, red-dotted line). Two wells were imaged for each pH-adjusted buffer tested, and three fields were acquired per well. The experiment was repeated three times. Geometrical objects indicate averaged ratios. SEM was within the geometrical objects. **(F-N)** Confocal microscopy micrographs of J774 macrophages loaded with dextrans labeled with ApHID (F-H), fluorescein (I-K), or Oregon Green (L-N) with Alexa 647 as a pH-independent dye. The cells were either fixed in 0.5% PFA and incubated in pH 5.0 buffer for 30 min at 37 °C (F, I, L) or imaged live at 37 °C in complete DMEM medium (G, J, M). In order to alkalinize acidic compartments, 20 mM methylamine was added to the cells (H, K, N). **(O-P)** Fluorescence ratios were interpolated to pH values using calibration curves prepared for each dye pair. To prepare the curves, J774 macrophages were fixed in 0.5% PFA and incubated in pH 5.0 buffer for 30 min and then imaged. Fluorescence ratios corresponding to pH 5.0 were calculated per LE/Ly or per field as described above and used to generate all subsequent ratios corresponding to pH values 3.5 and 5.0-7.4, using titration data obtained previously for the same dextrans in solution (see Suppl. Fig. S3 and Suppl. Table ST3). The resulting ratio-to-pH titrations were fit to 4-component sigmoidal curves. pH was calculated for each LE/Ly compartment (O) or field (P), and fields were averaged for each well imaged and condition (P). Two wells were imaged for each condition, and four fields were acquired for each well. The experiment was independently replicated three times. Geometrical objects and bars indicate averaged pH ±SD (O) or ±SEM (P). See Suppl. Table ST4 for statistics on pH measurements. Scale bars: 10 µm. Abbreviations: ‘OG’: Oregon Green; ‘A647’: Alexa Fluor 647; ‘ApH’: ApHID.

As expected, the fluorescein signal at pH 5.0 (Figs. 6I) or in living cells (Fig. 6J) was substantially weaker relative to that of ApHID (Figs. 6F-6G), whereas that of Oregon Green (Fig. 6L-6M) was comparable. When 20 mM methylamine was added to the media, ApHID signal decreased dramatically (Fig. 6H), but fluorescein and Oregon Green became substantially brighter (Fig. 6K, 6N). This is consistent with the pH-dependent fluorescence properties reported for these probes, which reflect the acidic nature of LE/Lys. Most importantly, these results indicate that, once in the LE/Lys, ApHID continues to be sensitive to changes in pH.

Next, we compared LE/Ly pH reported by each dye in living cells. pH-sensitive/Alexa 647 ratios were calculated and interpolated to pH for each imaged LE/Ly (Fig. 6O) or well (Fig. 6P, six wells imaged throughout three independent experiments) using a ratio-to-pH calibration curve for each probe. To prepare these curves, fluorescence ratios were determined for cells incubated in pH 5.0 buffer and thereafter used to generate all subsequent ratios (pH 3.5 and 5.0-7.4), using titration data in solution from the same dextrans. This approach is appropriate because the pH-dependent dynamic range of dextrans measured in cell culture is identical to that measured in solution (Fig. 6A-5E). It is important to note that in fixed cells, 50 mM TRIS maleate pH 5.0 buffer equilibrated well across cell membranes at 37 °C without causing significant membrane or cell swelling (Suppl. Fig. S1). However, membrane permeation was required for efficient buffer equilibration (Suppl. Fig. S2). Titration data obtained in solution was used to prepare the calibration curves for cell culture, and the data were fit to 4-component sigmoidal curves (Suppl. Fig. S3 and Suppl. Table ST3). The averaged interpolated lysosomal pH reported by ApHID after 1h incubation in DMEM at 37 °C was within 0.1 pH units of that reported by fluorescein and Oregon Green, and differences in pH between conditions were not statistically significant (Suppl. Table ST4).

Finally, in order to determine the stability of lysosomal pH over time, J774 macrophages with LE/Lys loaded with dextrans were equilibrated at 37 °C with 5% CO2 for 20 mins inside a confocal microscopy chamber and imaged every 5 minutes for 1h (see Methods section). Fluorescence ratios for each dye pair were interpolated to pH as described earlier and plotted as a function of time. Figure 7 shows LE/Ly pH over time reported by ApHID (Fig. 7A), fluorescein (Fig. 7B) or Oregon Green (Fig. 7C) (green). All probes reported stable pH at or near 5 with minimal variation over time. See Suppl. Table ST5 for statistics. These results indicate that LE/Ly pH measured using our ratiometric pH imaging methodology is stable irrespective of the pH-sensitive probe used as a reporter.

**Figure 7.**
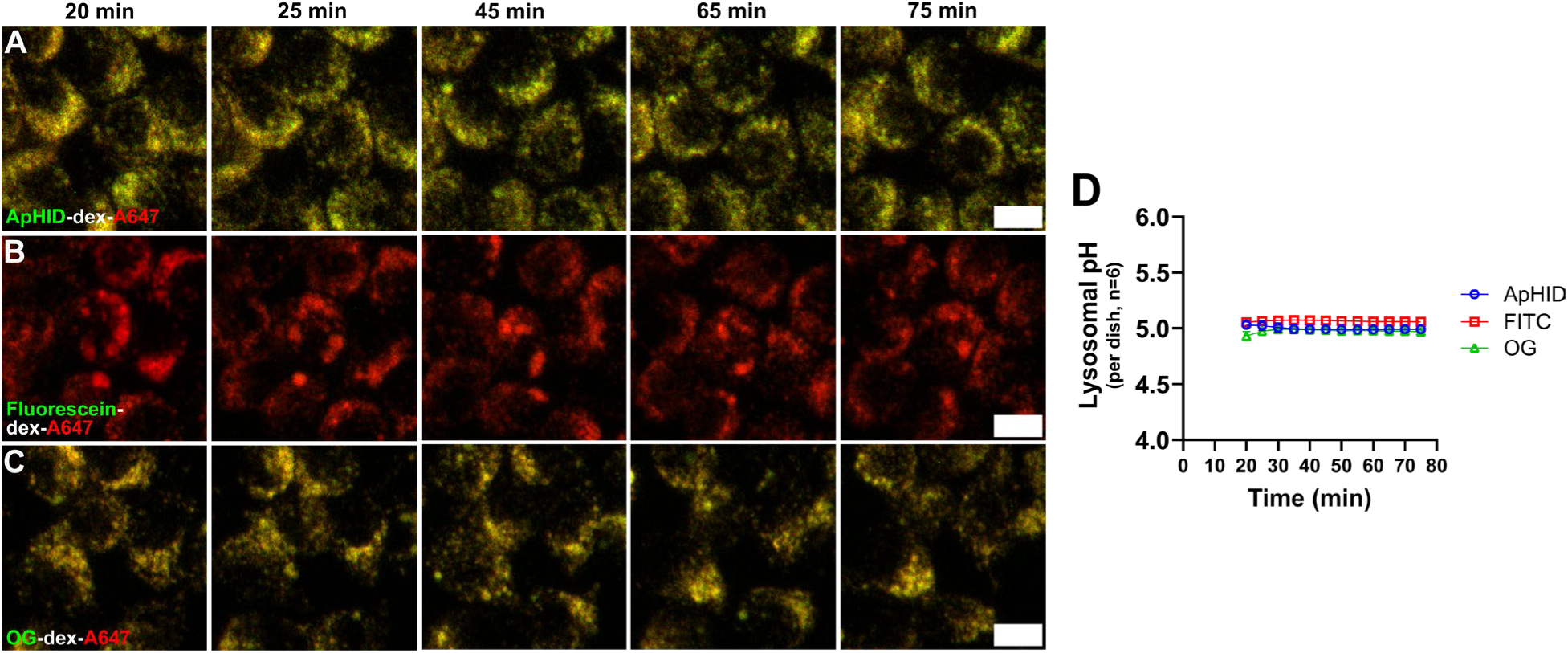
Lysosomal pH measured in J774 macrophages is stable over time. **(A-C)** J774 macrophage LE/Lys were loaded with dextrans labeled ApHID (A), fluorescein (B) or Oregon Green (C) and Alexa 647 (pH-independent) and imaged every 5 mins for 60 min using confocal microscopy. **(D)** pH-sensitive/Alexa 647 ratios measured per field were converted to pH values, and plotted against incubation time for each pH-sensitive probe. See Suppl. Table ST5 for statistics. Two wells were imaged per dye pair, 4 fields imaged per well. The experiment was replicated independently three times. Geometrical shapes and bars indicate average pH ±SEM. The SEM error bars fit within the symbols. Scale bars: 10 µm. Abbreviations: ‘OG’: Oregon Green; ‘A647’: Alexa Fluor 647.

In summary, ApHID reports virtually identical LE/Ly pH values as fluorescein and Oregon Green, and the reported LE/Ly acidity was found to be stable over time when imaged in equilibrated, constant temperature conditions. This validates the use of ApHID for ratiometric pH imaging of acidic vesicles in cell culture.

## DISCUSSION

We report the preparation and characterization of the novel pH-sensitive probe ApHID. The chemical design of ApHID confers the molecule with robust photostability and optimal spectroscopic properties to measure pH in the acidic range of endosomes and lysosomes. ApHID contains a PEG4 chain terminated in an *N*-hydroxysuccinimidyl ester reactive group, which increases solubility and allows for the derivatization of proteins and other molecules containing primary amines. In addition, ApHID is a weak base, whereas fluorescein, Oregon Green, and many other pH-sensitive probes are carboxylic acids, which is a novel feature. ApHID’s fluorescence increases with increasing acidity, whereas fluorescein and Oregon Green become dimmer. ApHID’s dynamic range between pH 4.0 and pH 6.0 was almost twice that of LysoSensor yellow/blue and Oregon Green, and four times greater than that of fluorescein. This allows for more accurate pH measurements in LE/Lys. Importantly, the pKa of ApHID and its pH-dependent fluorescence remain stable when exposed to high amounts of salt and protein as well as to •OH radicals at concentrations substantially higher than those found in endosomes and lysosomes. Accordingly, once endocytosed by J774 macrophages, ApHID fluorescence remained stable for an extended time. Altogether, the probe is robust to chemical modification, and it is much more photostable than the other pH indicator dyes we tested.

It is possible to derivatize amino-dextrans with various degrees of ApHID along with pH-independent fluorophores without altering ApHID’s fluorescence properties. This is advantageous when working with difficult imaging applications such as intravital imaging, which benefits from bright markers that can withstand sustained excitation. Interestingly, in derivatized dextrans, the pKa of ApHID can be modulated by altering the ionic environment on the polymers. We found that increasing dextran negative charge by derivatizing it with anionic molecules such as Alexa 405 increased ApHID’s pKa. This is most likely due to the stabilizing effect played by the nearby negative charge, which favors protonation of the aniline group in the ApHID core. By the same principle, adding positive charge to the polymer backbone should disfavor aniline protonation, thus requiring higher acidity to activate ApHID fluorescence, and ultimately lowering ApHID’s pKa. This demonstrates that ApHID pH sensitivity can be finely tuned by modulating amino-dextran net charge, which might be advantageous when pH sensing of different acidity ranges is required.

Cellular acidic compartments can be imaged live using nanosensors ^37^ or genetically encoded biosensors ^38,39^. The latter requires expressing the reporters in cell culture or in animal models, which is not trivial. However, dextrans labeled with pH-sensitive probes can be used to label acidic compartments in virtually any cell type. Labeling of polymers and loading and imaging of acidic compartments can be accomplished quickly within a couple of days. Furthermore, 70KDa dextrans remain inside LE/Lys for days and can be used to monitor pH for extended periods of time before being exocytosed. We demonstrate that ApHID can be used to measure the pH of acidic compartments in cultured cells. We validated our measurements by multiple comparisons with other pH-sensitive probes, fluorescein and Oregon Green, which are commonly used to track lysosomes and measure their lumenal pH. For ratiometric pH imaging, we derivatized dextrans with the various pH-sensitive probes and a second, pH-independent dye (Alexa 647) and loaded J774 macrophages with the polymers. Once endocytosed, dextrans reach LE/Lys, allowing for ratiometric imaging of the organelles. pH-sensitive/Alexa 647 ratios measured per field or for individual LE/Lys were interpolated to pH values using ratio-to-pH calibration curves. The accuracy of the ratio-to-pH calibration will determine the precision of the pH measurements. The pH distribution of individual LE/Lys reported by ApHID was tighter than that of Oregon Green, and similar to that of fluorescein. Differences between lysosomal pH measured with ApHID, fluorescein, and Oregon Green were minimal and not statistically significant. The pH reported by ApHID stayed within 0.1 pH units of that reported by the other dyes, indicating that they are effectively equivalent. As expected, once in acidic compartments, our probes were sensitive to changes in acidity induced by the addition of 20 mM methylamine to the DMEM medium, which alkalinized LE/Lys.

In summary, we believe that ApHID circumvents a number of limitations presented by most commercially available pH-sensitive probes. ApHID’s spectroscopic properties, pH-dependence between pH 4.0-6.0, as well as its resistance to oxidation and photobleaching, make it optimal for measuring LE/Ly pH in a variety of cell types, and we believe that our methodology will prove useful in demanding imaging applications such as intravital imaging of tissues.

## Supporting information

Supplementary Section

## RESOURCE AVAILABILITY

### Lead contact

Further information and requests for resources and reagents should be directed to and will be fulfilled by the lead contact, Dr. Santiago Sole-Domenech.

### Materials availability

Further information and requests for Acidic pH Indicator Dye (ApHID) should be directed to and will be fulfilled by the lead contact upon request.

### Data and code availability

Confocal microscopy stacks of images (Figures 5 and 6 and Suppl. Figs. 1 and 2) and all associated data analysis spreadsheets as well as all supporting supplementary data are stored at Weill Cornell Institutional Data Repository for Research (WIDRR) and will be made available by the lead contact upon request. This paper does not report original code. Any additional information required to reanalyze the data reported in this work paper is available from the lead contact upon request.

## ACKNOWLEDGEMENTS

This work was supported by the Cure Alzheimer’s Foundation grant CAF-211540-02 and NIH grants RF1-AG078244 and R01-HL093324. S.S.D. was supported by the Swedish Research Council International Postdoctoral Fellowship number 637-2013-503 / D0050301 and the Leon Levy Foundation Fellowship in Neuroscience. The authors are grateful to Weill Cornell Chemistry Core for synthesizing ApHID, Raksha Narendra for assistance with cell culture and dextran derivatization and Warren Zipfel at Cornell University for advice on quantum yield measurements.

## AUTHOR CONTRIBUTIONS

Conceptualization, F.R.M., S.S.D. J.D.W.; Methodology, S.S.D., F.R.M., P.K.S., J.D.W.; Validation, S.S.D., P.K.S.; Investigation, S.S.D., P.K.S.; Resources, F.R.M., S.S.D., J.D.W.; Data Curation, S.S.D., P.K.S.; Writing – Original Draft, S.S.D., F.R.M, J.D.W.; Writing – Review & Editing, S.S.D., F.R.M., J.D.W.; Visualization, S.S.D., Supervision, F.R.M., J.D.W.; Project Administration, F.R.M., S.S.D., J.D.W.; Funding Acquisition: F.R.M., S.S.D., J.D.W.

## METHODS

### Materials

A list of all reagents used and their source can be found in the Key Resource Table. Also, see Suppl. Table ST6 for a list of dextrans, concentrations, fluorophore labeling, and reaction conditions used for each experiment described below.

#### Dextran derivatization with fluorophores

Polymers of 10 KDa (Thermo Fisher D1860) or 70 KDa amino-dextrans (Thermo Fisher D1862 or Fina Biosolutions AD7×x33) were solubilized at various concentrations in sterile 0.1 M NaHCO3 buffer adjusted to pH 8.3 and reacted with the following *N*-hydroxysuccinimidyl esters (NHS): NHS-ApHID (custom-made), NHS-5/6-carboxyfluorescein (NHS-fluorescein, Thermo Fisher 46410), NHS-Oregon Green (Thermo Fisher O6147), NHS-Alexa 405 (Thermo Fisher A30000), NHS-Cy5•3xSO3^−^ (AstaTech 44193) or NHS-Alexa 647 (Thermo Fisher A20106) at various polymer:dye molar ratios (see Supplementary Section table ST3) for 1-2 h at room temperature (RT) with constant rotation. To examine the effect of various degrees of ApHID derivatization or the presence of negatively charged fluorophores on pKa we used 70 KDa dextrans from Fina Biosolutions, reacted with a fixed amount of NHS-Alexa 405 or NHS-Cy5•3xSO3^−^, followed by reaction with varying molar ratios of NHS-ApHID. For studies on the effect of charge density on ApHID pKa, the polymers were reacted with NHS-ApHID first, followed by aliquoting and reaction with various amounts of NHS-Alexa 405 to add negative charge density to the polymers. For ratiometric pH imaging experiments in live cells, we used 70 KDa dextrans from Thermo Fisher, reacted with NHS-ApHID, NHS-fluorescein or NHS-Oregon Green and NHS-Alexa 647 at 4:3 molar ratio. Following reaction, dextrans were purified by extensive dialysis in 3.5 KDa or 20 KDa cutoff Side-A-Lyzer dialysis cassettes (Thermo Fisher 66330 and 66003) against PBS.

#### Measuring labeling of dextrans with fluorophores by absorbance

When measuring ApHID incorporation, dextrans were diluted in 25 mM citric acid, 25 mM sodium citrate buffer adjusted to pH 3.0 and absorbance was read at 502 nm. When measuring labeling with fluorescein or Oregon Green, the dextrans were diluted in pH 7.4 PBS buffer, and absorbance was read at 490 nm. When measuring labeling with Alexa 405, Cy5•3xSO3^−^, or Alexa 647, the dextrans were diluted in buffer, and absorbance was read at 402, 666, or 655 nm, respectively. All dextrans were diluted to 0.02 mg/mL in buffers. Measurements were done using the quartz cuvette reader on a SpectraMax M3 plate reader (Molecular Devices). The extent of dextran labeling was lower than the polymer:dye ratio of the reaction mixture; dye incorporation efficiencies ranged from 25 to 50% (see Supplementary Section table ST3).

#### Buffers for measurements in solution

Dextrans were solubilized in buffers with pH adjusted between 1.5 and 8.5. To prepare the buffers, buffer salts were added to 0.66X PBS as follows: *pH 1.5 to 3.5 buffer:* 25 mM citric acid plus 25 mM sodium citrate; *pH 4.0 to 5.5:* 50 mM TRIS-maleate; *pH 6.0-7.0 buffer:* 50 mM sodium phosphate monobasic anhydrous; *pH 7.5:* 25 mM TRIS hydrochloride plus 25 mM sodium phosphate dibasic; *pH 8.0-8.5:* 50 mM TRIS base. The pH was determined using an Orion Star A211 pH meter (Thermo Fisher), and acidity was adjusted by adding 1-3N HCl or 1-10N NaOH dropwise. The resulting buffer ionic strength of the solutions was approximately 150 mM. After preparation, the solutions were filtered through a 0.4 µm filter membrane and stored at 4 °C.

#### Cell culture

J774A.1 murine macrophages (ATCC TIB-67) were grown in Dulbecco’s Modified Eagle’s Medium (DMEM) containing 4.5 g/L glucose and 1 mM sodium pyruvate (Corning 15-013-CV), with 10% fetal bovine serum (FBS, Gemini BenchMark FBS 100-106), 4 mM L-glutamine (Gibco 25030081), and 1% penicillin-streptomycin (Thermo Scientific 15140163) in an incubator at 37 °C with humidified atmosphere and 5% CO2. Cells were passed at a subcultivation ratio of 1:5 every 2-3 days.

#### Buffers and cell media for measurements in fixed and live cells

Fixed cells were incubated and imaged in buffers prepared by adding buffer salts to 0.13X PBS with 10% FBS, 40 mM methylamine hydrochloride, 40 mM sodium acetate, and 40 µM monensin (to ensure buffer equilibration across membranes) and with pH adjustment as described above. The resulting ionic strength of the solutions was approximately 150 mM. The solutions were filtered and stored at 4 °C. Live cells were imaged by confocal microscopy in a 96-well plate at 37 °C with 5% CO2, in DMEM without phenol red (Corning 90-013-PB) with 10% FBS, 2.2 g/L sodium bicarbonate (Sigma S5761), 4 mM L-glutamine, 1 mM sodium pyruvate (Sigma S8636) and 1% penicillin-streptomycin.

## Method Details

### pH-dependent absorbance measurements

To measure pH-dependent absorbance spectra for ApHID, 10 KDa amino-dextrans derivatized with the dye were diluted to 0.04 mg/mL in pH-adjusted buffers (see preparation of buffers above) and loaded into 384-well flat clear bottom black polystyrene microplates (Corning, 3746). Three wells were loaded per pH buffer condition and dye. The microplates were centrifuged at 3700 rpm for 1 min followed by absorbance spectra acquisition between 400-600 nm using a SpectraMax M3 plate reader. Absorbance was read from the top of the plate. To record pH-dependent absorbance titrations for all probes, derivatized 10 KDa amino-dextrans were diluted to 0.02 mg/mL in pH-adjusted buffers followed by absorbance reading at their excitation maxima of 500 nm (ApHID); 495 nm (fluorescein); 505 nm (Oregon Green); and 381 nm (LysoSensor yellow/blue). For both measurements, absorbance measured from buffers alone was subtracted as blank.

### pH-dependent fluorescence measurements

To record pH-dependent emission spectra for ApHID, 10 KDa amino-dextrans derivatized with the dye were diluted to 0.04 mg/mL in pH-adjusted buffers and loaded into 384-well microplates. ApHID was excited at 480 nm, and emission spectra were collected between 500 and 600 nm using a SpectraMax M3 plate reader. Fluorescence was read from the bottom of the plate. To record pH-dependent emission titrations, 10 KDa or 70 KDa amino-dextrans labeled with the various probes were diluted to 0.02 mg/mL in pH-adjusted buffers, followed by excitation and fluorescence measurement at their excitation and emission maxima (ApHID, ex/em 500/520 nm; fluorescein ex/em 490/525 nm; Oregon Green ex/em 505/525 nm; LysoSensor yellow/blue ex/em 381/521 nm; Alexa 405 ex/em 402/425 nm; Cy5•3xSO_3_ ^−^ ex/em 620/666 nm; Alexa 647 ex/em 620/655 nm). Emission measured from buffers alone was subtracted as blank. Measurements were repeated at least twice.

### Quantum yield and extinction coefficient measurements

ApHID quantum yield was determined following a previously published protocol ^40^. Stock solutions of NHS-ApHID and NHS-fluorescein were prepared in 1X PBS and allowed to hydrolyze overnight at RT. Four series of dilutions of ApHID, each in a different buffer, were prepared in buffers adjusted to pH 3.5 to 6.0. Four dilutions of fluorescein, with a known quantum yield of 0.95, were prepared in 0.1 M NaOH as a reference. For each dilution and buffer, absorbance and emission spectra were recorded in the range of 400-550 nm and 500-600 nm, respectively, using 1 cm quartz cuvettes and the cuvette reader in a SpectraMax M3 plate reader. Measurements in buffers alone were subtracted as blank. Integrated fluorescence was plotted against integrated absorbance for each dilution and buffer, and the curves were fit to a linear trend. Measurements were repeated twice. Quantum yields for ApHID, for each buffer pH, were calculated using the equation below:

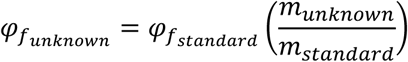

Where *munknown* and *mstandard* are the slopes of the resulting linear regression for ApHID and fluorescein dilutions, respectively.

To determine the extinction coefficient (ε) of ApHID, several dilutions of a hydrolyzed NHS-ApHID stock were prepared in pH 3.0 citric acid, sodium citrate buffer, and their absorbance at 492 nm was measured in quartz cuvettes using a SpectraMax M3 plate reader. Absorbance measurements in buffers alone were subtracted as blank. The corrected absorbance was fit to Beer’s law, A = ε·b·c, where A is the absorbance, ε is the molar extinction coefficient, b is the path length of the cuvette and c is the concentration. Measurements were repeated twice.

### Hydroxyl radical (·OH) generation and oxidation assay

500 µM NHS-ApHID, NHS-fluorescein, and NHS-Oregon Green solutions were prepared in 1X PBS and allowed to hydrolyze overnight. The probes were then diluted into 10 µM aliquots in 1X PBS and refrigerated in ice. Stock solutions of 50 mM Fe(OCl^-^)2 (Sigma-Aldrich 334081) and 50 mM H2O2 (VWR, 30% aqueous stock, BDH7690-3) were prepared in ultra-pure water and immediately added to the fluorophore aliquots to a final concentration of 100 or 200 µM. Plastic tubes containing the aliquots were vortexed for 10 seconds, sealed in paraffin, and incubated at 37 °C in a convection oven with constant rotation for 20h. Following reaction, the aliquots were diluted 1:200 in buffers with pH adjusted between 3.5 and 7.5 and loaded into 384-well microplates. Fluorescence was measured for each probe and buffer at their ex/em maxima using a SpectraMax M3 plate reader. The experiment was repeated twice.

### Effect of salt and protein on probe fluorescence and pKa

10 KDa amino-dextrans derivatized with the various probes were diluted to 0.02 mg/mL in pH-adjusted buffers additionally supplemented with either 1 mM CaCl2 and 1 mM MgCl2, sodium acetate (replacing sodium chloride), or 50 mg/mL bovine serum albumin (BSA). pH-adjusted buffers lacking sodium chloride were prepared using a 0.66X PBS base for which sodium chloride had been replaced by sodium acetate. Buffers containing 50 mg/mL BSA were prepared in 1X PBS and adjusted to pH 2.5-7.0. Dextran dilutions in buffers were vortexed and loaded into 384-well microplates. The microplates were sealed using paraffin and incubated at 37 °C in a convection oven for 20h. After incubation, the fluorescence of the various probes was measured at their ex/em maxima using a SpectraMax M3 plate reader. The experiment was repeated twice.

### Photobleaching study in fixed J774 macrophages using confocal microscopy and digital image analysis

J774A.1 murine macrophages were seeded in 35-mm dishes with central 7 mm diameter glass bottom imaging chambers coated with poly-D lysine at 10,000 cells per chamber and allowed to settle for 1-2 h in an incubator at 37 °C. Once settled, cells were incubated with 70 KDa dextrans from Thermo Fisher previously labeled with NHS-ApHID, NHS-fluorescein, or NHS-Oregon Green at a final concentration of 1 mg/mL in complete DMEM overnight. The following morning, cells were washed once and chased for 3h in fresh DMEM, followed by 5 min fixation in 0.5% paraformaldehyde (PFA) and three washes in 1X PBS. Immediately prior to imaging, cells labeled with ApHID or Oregon Green dextrans were incubated in TRIS maleate pH 5.0 buffer, and cells labeled with fluorescein dextran were incubated in pH 7.4 1X PBS buffer, for 5 min at 37 °C inside the confocal microscope incubation chamber. The buffers were supplemented with 40 mM methylamine hydrochloride, 40 mM sodium acetate, 2.5 µM nigericin, and 2.5 µM monensin to facilitate buffer equilibration across cell membranes. Following buffer equilibration, cells were immediately imaged by confocal microscopy. 2 dishes per fluorophore condition were imaged, and 4 fields were acquired for each dish. The experiment was independently replicated twice.

### Ratiometric imaging of fixed J774 macrophages in pH 4.0 to pH 6.0 buffers at 27 °C

J774A.1 murine macrophages were seeded in 96-well plates with transparent polymer bottoms (Cellvis, P96-1.5P) at 40,000 cells per well and allowed to settle for 1-2 h in the incubator at 37 °C. Once settled, cells were incubated with 70 KDa amino-dextrans from Thermo Fisher labeled with NHS-ApHID (pH-sensitive) and NHS-Alexa 647 (pH-independent) at a 4:3 molar ratio, with a final dextran concentration of 0.5 mg/mL in complete DMEM overnight. The following morning, cells were washed twice and chased in fresh DMEM for at least 3h. Next, the cells were fixed in 0.5% PFA for 5 mins and washed 3x in 1X PBS, and the plate was then transferred to the confocal microscope incubator and allowed to equilibrate at 27 °C for at least 20 mins followed by image acquisition (time 0). After that, the PBS buffer was carefully removed from the wells using a 200 µL pipette followed by the addition of 200 µL of buffers with pH adjusted to 4.0, 4.5, 5.0, 5.5, and 6.0 (see main Methods section) supplemented with 10% FBS, 40 mM methylamine, 40 mM sodium acetate, and 40 µM monensin (buffers had been pre-warmed to 27 °C prior to addition to cells). The cells were allowed to equilibrate in the buffers for 5 mins, and stacks of images were then acquired every 5 mins for a period of 2h. 2 wells were imaged for each buffer condition, and 3 fields were acquired per well. The experiment was independently replicated three times. ApHID/Alexa 647 ratios were calculated as described in the Methods section for ratiometric pH imaging and plotted against incubation time for each buffer condition. Data were fit to a rectangular hyperbola for visualization purposes.

### Ratiometric imaging in TRIS maleate pH 5.0 buffer containing various amounts of monensin at 37 °C

Cells were seeded in 96-well plates, labeled with ApHID-Alexa647 dextrans, and fixed in PFA as described above. The plate was then transferred to the confocal inset and allowed to equilibrate at 37 °C for at least 20 mins followed by image acquisition (time 0). After that, the PBS buffer was carefully removed from the wells using a 200 µL pipette followed by the addition of 200 µL of 50 mM TRIS maleate pH 5.0 buffer supplemented with 10% FBS, 40 mM methylamine, 40 mM sodium acetate, and 5, 40, or 80 µM monensin, pre-warmed at 37 °C prior to addition. The cells were allowed to equilibrate in the buffers for 6 mins, and stacks of images were acquired every 5 mins for a period of 40 min. Two wells were imaged for each buffer condition, and 3 fields were acquired per well. The experiment was independently replicated three times. ApHID/Alexa 647 ratios were calculated as described in the Methods section for ratiometric pH imaging and plotted against incubation time for each buffer condition. Data were fit to a rectangular hyperbola for visualization purposes only.

### Ratiometric pH calibration of fixed J774 macrophage lysosomes

J774A.1 murine macrophages were seeded in 96-well plates with glass-like polymer bottom (Cellvis, P96-1.5P) at 40,000 cells per well and allowed to settle for 1-2 h in the incubator at 37 °C. Once settled, cells were incubated with 70 KDa amino-dextrans from Thermo Fisher labeled with NHS-ApHID (pH-sensitive) and NHS-Alexa 647 (pH-independent) at a 4:3 molar ratio, with a final concentration of 0.5 mg/mL in complete DMEM overnight. The following morning, cells were washed twice and chased in fresh DMEM for at least 3h. Next, the cells were fixed in 0.5% PFA for 5 mins and washed 3x in 1X PBS, and the plate was then transferred to an incubation chamber and allowed to equilibrate to 37 °C for at least 20 mins, followed by image acquisition (time 0). After that, the PBS buffer was carefully removed from the wells using a 200 µL pipette and 200 µL of buffers with pH adjusted to 4.0, 4.5, 5.0, 5.5, and 6.0 (see main Methods section) were added. The buffers were supplemented with 10% FBS, 40 mM methylamine, 40 mM sodium acetate and 40 µM monensin and had been pre-warmed to 37 °C prior to addition to cells. The cells were allowed to equilibrate in the buffers for 20 min (pH 4.0 and pH 4.5) or at least 30 min (pH 5.0-6.0) followed by image stack acquisition. Two wells were imaged for each buffer condition, and 3 fields were acquired per well. The experiment was independently replicated three times.

### Live ratiometric pH imaging of J774 macrophage lysosomes

J774A.1 murine macrophages were seeded in 96-well plates with glass-like polymer bottom (Cellvis, P96-1.5P) at 40,000 cells per well and allowed to settle for 1-2 h in the incubator at 37 °C. Once settled, cells were incubated with 70 KDa amino-dextrans from Thermo Fisher labeled with NHS-ApHID, NHS-fluorescein or NHS-Oregon Green (pH-sensitive), and NHS-Alexa 647 (pH-independent) at a 4:3 molar ratio, with a final concentration of 0.5 mg/mL in complete DMEM overnight. The following morning, cells were washed twice and chased in fresh DMEM for at least 3h, followed by 1h equilibration in imaging DMEM without phenol red. The plate was then transferred to the confocal microscope incubation chamber and allowed to equilibrate at 37 °C with 5% CO2 for 20 min prior to imaging. Cells were imaged in DMEM without phenol red. 20 mM methylamine hydrochloride was added to some wells and incubated for 10 min. Stacks of images were acquired from each well every 5 mins, for 1h. For pH calibration, two wells for each fluorophore pair were fixed in 0.5% PFA for 5 min and washed three times in 1X PBS, followed by incubation in 50 mM TRIS maleate pH 5.0 buffer supplemented with 10% FBS, 40 mM methylamine hydrochloride, 40 mM sodium acetate and 40 µM monensin, at 37 °C for 30 min inside the confocal microscope incubation chamber, and imaged immediately after. Two wells were imaged for each condition and dye pair, and four fields were acquired for each well. The experiment was independently replicated three times.

### Confocal microscopy

#### Photobleaching study in fixed J774 macrophages

J774 macrophages were imaged with an LSM 880 confocal microscope (Zeiss) at 37 °C. Fluorophores were excited using a 35 mW 488 nm laser with a digital power output adjusted to 30%, yielding 5.2 µW of power at the front element of the 40X objective (1.30 NA) used for imaging, as determined with a power meter. Single cell planes were irradiated for 0.5 seconds per cycle (50 cycles in total) with 1 second intervals between irradiation pulses. Fluorescence was detected using a high-sensitivity Zeiss GaAs detector with a spectral window adjusted to collect light at 500-550 nm. Pixel dwell time was 0.33 µs.

#### Ratiometric pH imaging of J774 macrophage lysosomes

Imaging of fixed cells or live cells was done with a Leica Stellaris confocal microscope using a 40X air objective (0.95 NA) with pinhole adjusted to 1 Airy unit. ApHID, fluorescein and Oregon Green were excited using a white light solid-state laser adjusted to 495 nm and Alexa 647 was excited with the laser adjusted to 650 nm. Fluorescence was detected using high-sensitivity silicon-based HyD detectors with a spectral window adjusted to collect light between 500-550 nm and 660-720 nm for all green-emitting dyes and Alexa 647, respectively. Stacks of images with 1.5 µm separation in the vertical axis were acquired for each field imaged.

#### Ratiometric imaging of buffer equilibration assays

Imaging was done with a Leica Stellaris confocal microscope using a 40X air objective (0.95 NA) with the pinhole adjusted to 1 Airy unit. ApHID was excited using a white light solid-state laser adjusted to 495 nm, and Alexa 647 was excited with the laser adjusted to 650 nm. Fluorescence was detected using high-sensitivity silicon-based HyD detectors with a spectral window adjusted to collect light between 500-550 nm and 660-720 nm for all green-emitting dyes and Alexa 647, respectively. Stacks of images with 1.5 µm separation in the vertical axis were acquired for each field imaged.

### Digital image analysis

Digital image analysis and quantification was done using FIJI (ImageJ) ^41^ version 1.54f for Windows (https://fiji.sc/) and MetaMorph version 6.7.1 for Windows (Molecular Devices, San Jose, California USA, www.moleculardevices.com).

#### Photobleaching study in fixed J774 macrophages

Stacks of images for each acquired field were corrected for background intensity by subtracting the 5^th^ percentile intensity value for each image in the stack. To quantify fluorescence signal per field, a sum projection for each stack (sum of each image in the stack) was generated, and total integrated intensity (F) per field was measured. Fluorescence was normalized to the intensity corresponding to the first irradiation cycle (F0) and plotted as F/F0 against cycle time.

#### Ratiometric pH imaging of J774 macrophage lysosomes

Stacks of images for each acquired field were corrected for background intensity by subtracting the 5^th^ percentile intensity value for each image in the stack. To quantify pH per field, a sum projection for each was generated, and fluorescence signal from organelles was selected using an intensity threshold applied to the pH-independent channel (Alexa 647). A mask was then generated and transferred to the pH-dependent channel. Integrated intensity was measured for each masked channel, and pH-sensitive/pH-independent ratios were calculated for each field. To quantify pH per LE/Ly, the fluorescence signal from organelles was selected using an intensity threshold applied to the pH-independent channel, for each individual plane. The internally thresholded objects function in MetaMorph was used to identify individual objects corresponding to labeled organelles. Two adjacent objects would be separated when the peak intensities of their Gaussian distribution differed by at least 50%. Dissected objects were analyzed using the integrated morphometry analysis function with a circularity filter set to 90%. Finally, integrated intensity of the pH-sensitive and pH-independent channels was measured for each dissected object, and their ratio values were calculated. Ratios calculated for each field or object imaged were interpolated to pH values using ratio-to-pH calibration curves. To prepare the curves, fixed J774 macrophages loaded with derivatized dextrans were incubated in 50 mM TRIS maleate pH 5.0 buffer and imaged as described earlier. The pH 5.0 ratios were determined per field or per object and used to generate pH-dependent ratio values corresponding to pH 3.5-7.4 buffers, using titration data for the same dextrans in solution. The resulting ratio-to-pH calibrations were fit to 4-component sigmoidal curves.

### Statistical data analysis

Statistical analyses were performed using GraphPad Prism version 10.3.1 for Windows (GraphPad Software, Boston, Massachusetts, USA, www.graphpad.com). Experiments in solution were repeated independently at least twice, and averages ± SEM are shown (Figs. 1, 2 and 4 and Suppl. Fig. S3). Photobleaching measurements in fixed cells were repeated twice; two dishes were imaged per condition and experiment, and 4 fields were acquired per dish. Averaged fluorescence intensity per irradiation cycle per field (16 fields in total) ± SEM is presented (Fig. 3). For buffer equilibration assays, experiments were repeated three times; 2 wells were imaged per buffer condition and 3 fields were acquired per well (Suppl. Figs. S1 and S2). For ratiometric pH imaging in fixed or live cells, experiments were repeated three times; two wells were imaged per condition and probe (6 wells in total), and three to four fields were acquired per well (Fig. 5 and 6). Averaged pH per well ±SEM or averaged pH per LE/Ly ±SD are presented. Differences between averaged LE/Ly pH per well reported by the various probes and between conditions were assessed using the unpaired One-way ANOVA followed by Dunnett’s multiple comparison test (Fig. 5). P-values are shown as p>0.05 (ns), p≤0.05 (*), p≤0.01 (**), p≤0.001 (***), and p≤0.0001 (****). Brown-Forsythe’s and Bartlett’s tests were used to assess differences in data’s standard deviations between conditions. Interval confidence was set to 95%.

## REFERENCES

1. Mukherjee, S., Ghosh, R.N., and Maxfield, F.R. (1997). Endocytosis. Physiol Rev 77, 759–803. 10.1152/physrev.1997.77.3.759.

2. Lubke, T., Lobel, P., and Sleat, D.E. (2009). Proteomics of the lysosome. Biochim Biophys Acta 1793, 625–635. 10.1016/j.bbamcr.2008.09.018.

3. Sole-Domenech, S., Rojas, A.V., Maisuradze, G.G., Scheraga, H.A., Lobel, P., and Maxfield, F.R. (2018). Lysosomal enzyme tripeptidyl peptidase 1 destabilizes fibrillar Abeta by multiple endoproteolytic cleavages within the beta-sheet domain. Proc Natl Acad Sci U S A 115, 1493–1498. 10.1073/pnas.1719808115.

4. Yamashiro, D.J., and Maxfield, F.R. (1987). Kinetics of endosome acidification in mutant and wild-type Chinese hamster ovary cells. J Cell Biol 105, 2713–2721. 10.1083/jcb.105.6.2713.

5. Maxson, M.E., and Grinstein, S. (2014). The vacuolar-type H(+)-ATPase at a glance - more than a proton pump. J Cell Sci 127, 4987–4993. 10.1242/jcs.158550.

6. Maxfield, F.R. (2014). Role of endosomes and lysosomes in human disease. Cold Spring Harb Perspect Biol 6, a016931. 10.1101/cshperspect.a016931.

7. Aits, S., and Jaattela, M. (2013). Lysosomal cell death at a glance. J Cell Sci 126, 1905–1912. 10.1242/jcs.091181.

8. Halle, A., Hornung, V., Petzold, G.C., Stewart, C.R., Monks, B.G., Reinheckel, T., Fitzgerald, K.A., Latz, E., Moore, K.J., and Golenbock, D.T. (2008). The NALP3 inflammasome is involved in the innate immune response to amyloid-beta. Nat Immunol 9, 857–865. 10.1038/ni.1636.

9. Lee, J.H., Yang, D.S., Goulbourne, C.N., Im, E., Stavrides, P., Pensalfini, A., Chan, H., Bouchet-Marquis, C., Bleiwas, C., Berg, M.J., et al. (2022). Faulty autolysosome acidification in Alzheimer’s disease mouse models induces autophagic build-up of Aβ in neurons, yielding senile plaques. Nat Neurosci 25, 688–701. 10.1038/s41593-022-01084-8.

10. Colacurcio, D.J., and Nixon, R.A. (2016). Disorders of lysosomal acidification-The emerging role of v-ATPase in aging and neurodegenerative disease. Ageing Res Rev 32, 75–88. 10.1016/j.arr.2016.05.004.

11. Majumdar, A., Cruz, D., Asamoah, N., Buxbaum, A., Sohar, I., Lobel, P., and Maxfield, F.R. (2007). Activation of microglia acidifies lysosomes and leads to degradation of Alzheimer amyloid fibrils. Mol Biol Cell 18, 1490–1496. 10.1091/mbc.e06-10-0975.

12. Sole-Domenech, S., Cruz, D.L., Capetillo-Zarate, E., and Maxfield, F.R. (2016). The endocytic pathway in microglia during health, aging and Alzheimer’s disease. Ageing Res Rev 32, 89–103. 10.1016/j.arr.2016.07.002.

13. Maxfield, F.R., and McGraw, T.E. (2004). Endocytic recycling. Nat Rev Mol Cell Biol 5, 121–132. 10.1038/nrm1315.

14. Ohkuma, S., and Poole, B. (1978). Fluorescence probe measurement of the intralysosomal pH in living cells and the perturbation of pH by various agents. Proc Natl Acad Sci U S A 75, 3327–3331. 10.1073/pnas.75.7.3327.

15. Feng, T., Sheng, R.R., Sole-Domenech, S., Ullah, M., Zhou, X., Mendoza, C.S., Enriquez, L.C.M., Katz, II, Paushter, D.H., Sullivan, P.M., et al. (2020). A role of the frontotemporal lobar degeneration risk factor TMEM106B in myelination. Brain 143, 2255–2271. 10.1093/brain/awaa154.

16. Majumdar, A., Capetillo-Zarate, E., Cruz, D., Gouras, G.K., and Maxfield, F.R. (2011). Degradation of Alzheimer’s amyloid fibrils by microglia requires delivery of ClC-7 to lysosomes. Mol Biol Cell 22, 1664–1676. 10.1091/mbc.E10-09-0745.

17. Canton, J., and Grinstein, S. (2015). Measuring lysosomal pH by fluorescence microscopy. Methods Cell Biol 126, 85–99. 10.1016/bs.mcb.2014.10.021.

18. Johnson, D.E., Ostrowski, P., Jaumouille, V., and Grinstein, S. (2016). The position of lysosomes within the cell determines their luminal pH. J Cell Biol 212, 677–692. 10.1083/jcb.201507112.

19. Haggie, P.M., and Verkman, A.S. (2007). Cystic fibrosis transmembrane conductance regulator-independent phagosomal acidification in macrophages. J Biol Chem 282, 31422–31428. 10.1074/jbc.M705296200.

20. Haggie, P.M., and Verkman, A.S. (2009). Unimpaired lysosomal acidification in respiratory epithelial cells in cystic fibrosis. J Biol Chem 284, 7681–7686. 10.1074/jbc.M809161200.

21. Zen, K., Biwersi, J., Periasamy, N., and Verkman, A.S. (1992). Second messengers regulate endosomal acidification in Swiss 3T3 fibroblasts. J Cell Biol 119, 99–110. 10.1083/jcb.119.1.99.

22. Bayer, N., Schober, D., Prchla, E., Murphy, R.F., Blaas, D., and Fuchs, R. (1998). Effect of bafilomycin A1 and nocodazole on endocytic transport in HeLa cells: implications for viral uncoating and infection. J Virol 72, 9645–9655. 10.1128/JVI.72.12.9645-9655.1998.

23. Cain, C.C., Sipe, D.M., and Murphy, R.F. (1989). Regulation of endocytic pH by the Na+,K+-ATPase in living cells. Proc Natl Acad Sci U S A 86, 544–548. 10.1073/pnas.86.2.544.

24. Lee, J.H., Yu, W.H., Kumar, A., Lee, S., Mohan, P.S., Peterhoff, C.M., Wolfe, D.M., Martinez-Vicente, M., Massey, A.C., Sovak, G., et al. (2010). Lysosomal proteolysis and autophagy require presenilin 1 and are disrupted by Alzheimer-related PS1 mutations. Cell 141, 1146–1158. 10.1016/j.cell.2010.05.008.

25. Lee, J.H., Wolfe, D.M., Darji, S., McBrayer, M.K., Colacurcio, D.J., Kumar, A., Stavrides, P., Mohan, P.S., and Nixon, R.A. (2020). beta2-adrenergic Agonists Rescue Lysosome Acidification and Function in PSEN1 Deficiency by Reversing Defective ER-to-lysosome Delivery of ClC-7. J Mol Biol 432, 2633–2650. 10.1016/j.jmb.2020.02.021.

26. Haka, A.S., Grosheva, I., Chiang, E., Buxbaum, A.R., Baird, B.A., Pierini, L.M., and Maxfield, F.R. (2009). Macrophages create an acidic extracellular hydrolytic compartment to digest aggregated lipoproteins. Mol Biol Cell 20, 4932–4940. 10.1091/mbc.e09-07-0559.

27. Jacquet, R.G., Gonzalez Ibanez, F., Picard, K., Funes, L., Khakpour, M., Gouras, G.K., Tremblay, M.E., Maxfield, F.R., and Sole-Domenech, S. (2024). Microglia degrade Alzheimer’s amyloid-beta deposits extracellularly via digestive exophagy. Cell Rep 43, 115052. 10.1016/j.celrep.2024.115052.

28. Maeda, H., Kowada, T., Kikuta, J., Furuya, M., Shirazaki, M., Mizukami, S., Ishii, M., and Kikuchi, K. (2016). Real-time intravital imaging of pH variation associated with osteoclast activity. Nat Chem Biol 12, 579–585. 10.1038/nchembio.2096.

29. Soares, M.P., and Hamza, I. (2016). Macrophages and Iron Metabolism. Immunity 44, 492–504. 10.1016/j.immuni.2016.02.016.

30. Collin, F. (2019). Chemical Basis of Reactive Oxygen Species Reactivity and Involvement in Neurodegenerative Diseases. Int J Mol Sci 20. 10.3390/ijms20102407.

31. Lyublinskaya, O., and Antunes, F. (2019). Measuring intracellular concentration of hydrogen peroxide with the use of genetically encoded H(2)O(2) biosensor HyPer. Redox Biol 24, 101200. 10.1016/j.redox.2019.101200.

32. Sies, H. (2017). Hydrogen peroxide as a central redox signaling molecule in physiological oxidative stress: Oxidative eustress. Redox Biol 11, 613–619. 10.1016/j.redox.2016.12.035.

33. Vaneev, A.N., Gorelkin, P.V., Garanina, A.S., Lopatukhina, H.V., Vodopyanov, S.S., Alova, A.V., Ryabaya, O.O., Akasov, R.A., Zhang, Y., Novak, P., et al. (2020). In Vitro and In Vivo Electrochemical Measurement of Reactive Oxygen Species After Treatment with Anticancer Drugs. Anal Chem 92, 8010–8014. 10.1021/acs.analchem.0c01256.

34. Christensen, K.A., Myers, J.T., and Swanson, J.A. (2002). pH-dependent regulation of lysosomal calcium in macrophages. J Cell Sci 115, 599–607. 10.1242/jcs.115.3.599.

35. Murphy, E. (2000). Mysteries of magnesium homeostasis. Circ Res 86, 245–248. 10.1161/01.res.86.3.245.

36. Hoffmann, B., and Kosegarten, H. (1995). FITC-dextran for measuring apoplast pH and apoplastic pH gradients between various cell types in sunflower leaves. Physiologia Plantarum 95, 327–335. 10.1111/j.1399-3054.1995.tb00846.x.

37. Kim, M., Chen, C., Yaari, Z., Frederiksen, R., Randall, E., Wollowitz, J., Cupo, C., Wu, X., Shah, J., Worroll, D., et al. (2023). Nanosensor-based monitoring of autophagy-associated lysosomal acidification in vivo. Nat Chem Biol 19, 1448–1457. 10.1038/s41589-023-01364-9.

38. Ponsford, A.H., Ryan, T.A., Raimondi, A., Cocucci, E., Wycislo, S.A., Frohlich, F., Swan, L.E., and Stagi, M. (2021). Live imaging of intra-lysosome pH in cell lines and primary neuronal culture using a novel genetically encoded biosensor. Autophagy 17, 1500–1518. 10.1080/15548627.2020.1771858.

39. Lee, J.H., Rao, M.V., Yang, D.S., Stavrides, P., Im, E., Pensalfini, A., Huo, C., Sarkar, P., Yoshimori, T., and Nixon, R.A. (2019). Transgenic expression of a ratiometric autophagy probe specifically in neurons enables the interrogation of brain autophagy in vivo. Autophagy 15, 543–557. 10.1080/15548627.2018.1528812.

40. Magde, D., Wong, R., and Seybold, P.G. (2002). Fluorescence quantum yields and their relation to lifetimes of rhodamine 6G and fluorescein in nine solvents: improved absolute standards for quantum yields. Photochem Photobiol 75, 327–334. 10.1562/0031-8655(2002)075<0327:fqyatr>2.0.co;2.

41. Schindelin, J., Arganda-Carreras, I., Frise, E., Kaynig, V., Longair, M., Pietzsch, T., Preibisch, S., Rueden, C., Saalfeld, S., Schmid, B., et al. (2012). Fiji: an open-source platform for biological-image analysis. Nat Methods 9, 676–682. 10.1038/nmeth.2019.

