## Supplementary Section for "Real-Time Ratiometric pH Imaging of Macrophage Lysosomes Using the Novel pH-sensitive Probe ApHID"

**1. Rationale behind the use of 50 mM TRIS maleate pH 5.0 buffer to incubate fixed cells for a fluorescence ratio-to-pH calibration (supporting main section Figure 6 and 7).**

*a) Buffers containing membrane-permeant equilibrators adjust lysosomal pH within minutes.*

To generate ratio-to-pH calibration curves, we measure pH-sensitive/pH-independent ratios in lysosomes of fixed cells, labeled with fluorescent dextrans. We incubate the cells with 50 mM TRIS maleate pH 5.0 buffer supplemented with membrane-permeant pH equilibrators (sodium acetate, methylamine hydrochloride, and monensin) [1,2]. This yields a ratio that corresponds to pH 5.0 in cells. This ratio value can be used to generate all other ratios of the calibration curve (corresponding to pH 3.5-4.5 and 5.5-7.4). To do so, we use titration data obtained in solution. This approach is possible because the pH-dependent fluorescence ratio of ApHID-Alex647 dextrans measured in cell culture, using confocal microscopy, is identical to that measured in solution (see main section, Figure 6A-6E). Also, the pH-dependence of ApHID remains constant within a wide range of labeling of the dextran polymer (see main section, Figure 4). Therefore, dextran batches with different degrees of dye incorporation still show the same pH response.

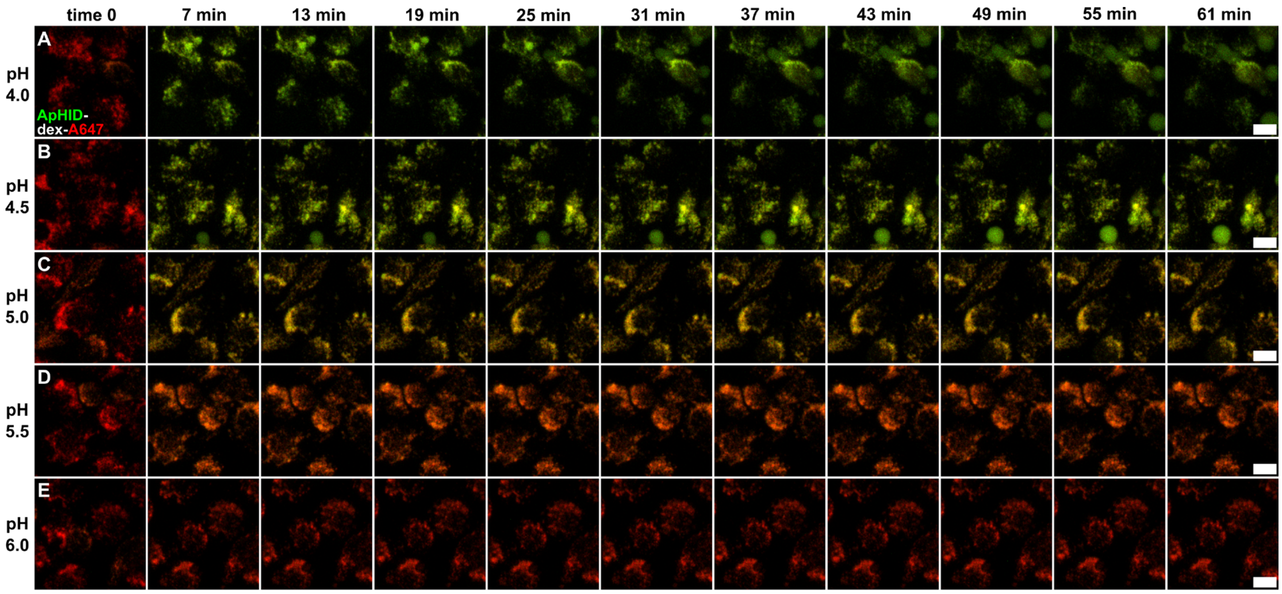


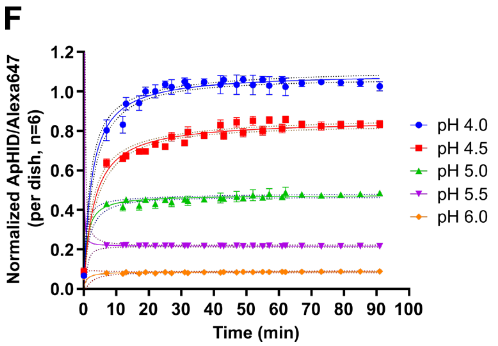


**Suppl. Fig S1. Time course of pH equilibration in buffers containing membrane-permeant equilibrators in J774 macrophages at 27 ºC. (A-E)** J774 macrophage lysosomes were loaded with dextrans labeled with ApHID and Alexa 647 (pH-independent) and fixed with 0.5% PFA. The cells were then incubated in 1X PBS (time 0, A-E) and then with various pH-adjusted buffers (pH 4.0 to 6.0) for 1h and imaged every 6 mins using confocal microscopy. The buffers contained 40 mM methylamine, 40 mM sodium acetate 40 µM monensin as membrane-permeant pH equilibrators. Overlays showing ApHID (green) and Alexa 647 (red) channels are shown for each buffer condition. **(F)** ApHID/Alexa 647 ratios measured for each buffer condition were normalized to pH 4.0, 20 min time point and plotted against incubation time. Data were fit to a rectangular hyperbola for visualization purposes only. Two wells per buffer pH condition were imaged, and 3 fields were acquired per well. The experiment was independently replicated three times. Geometrical objects and bars indicate averages ±SEM. Scale bars: 10 µm. Abbreviations: 'A647': Alexa Fluor 647.

Supp. Fig. S1 shows fluorescence measurements of ApHID (green) and Alexa 647 (red) in fixed J774 macrophages with lysosomes labeled with ApHID-Alexa647-dextran. The cells were incubated with various pH-adjusted buffers (see Methods section). All buffers contained 40 mM methylamine, 40 mM sodium acetate and 40 µM monensin as membrane-permeant equilibrators, and the cells were imaged at 27 ºC. Prior to buffer addition, cells in 1X PBS (pH 7.4) showed almost no ApHID signal (S1A-S1E, time 0). When buffers were added, ApHID signal increased over time, proportionally to buffer acidity (S1A-S1E). As expected, ApHID signal was brightest at pH 4.0.

However, at that pH, some cells started to show membrane swelling and cytosolic dextran fluorescence (S1A). This is likely due to lysosomal membrane damage and/or permeabilization, which caused dextran leakage. This effect was also seen, although to a lesser extent, when pH 4.5 buffer was used (S1B). Cells incubated in pH 5.0-6.0 buffers did not show any lysosomal membrane damage or fluorescence leakage over time (S1C-S1E). ApHID/Alexa 647 ratio equilibration reached a stable plateau in pH 5.0-6.0 buffers within 30 mins of incubation, whereas equilibration was slower in pH 4.0 and pH 4.5 buffers (S1A-S1E, S1F). Based on that, the use of 50 mM TRIS maleate pH 5.0 buffer for our calibrations seems optimal.

*b) Supplementation of pH 5.0 TRIS maleate buffer with monensin facilitates buffer equilibration across membranes at 37 ºC.*

Fixed cells used to prepare ratio-to-pH calibrations are incubated with pH 5.0 buffer supplemented with the *membrane-permeant equilibrators* methylamine, sodium acetate, and monensin, in order to facilitate buffer diffusion across cell membranes. Using high concentrations of monensin, however, can lead to membrane permeabilization and dextran leakage, yielding inaccurate pH-sensitive/pH-independent ratios and erroneous calibration curves. To determine the optimal monensin concentration to use, we incubated fixed J774 macrophages, loaded with ApHID-Alexa647 dextrans, with 50 mM TRIS maleate pH 5.0 buffer (see earlier section) supplemented with 0, 5, 40, or 80 µM monensin, and measured ApHID/Alexa647 ratio at 37 ºC over time.

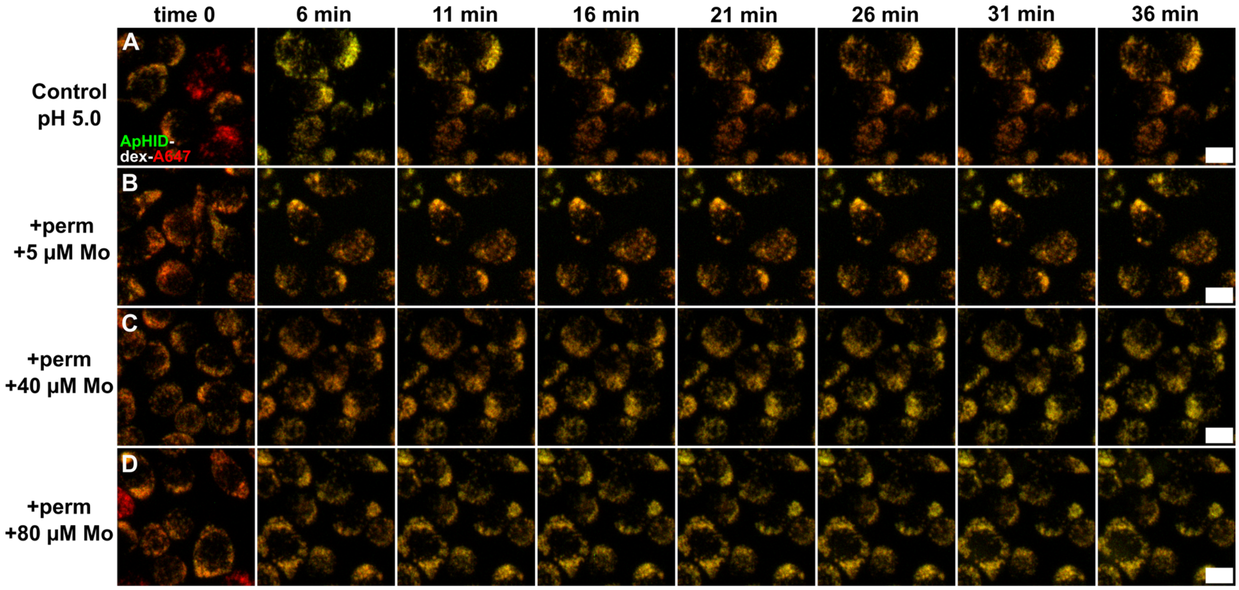


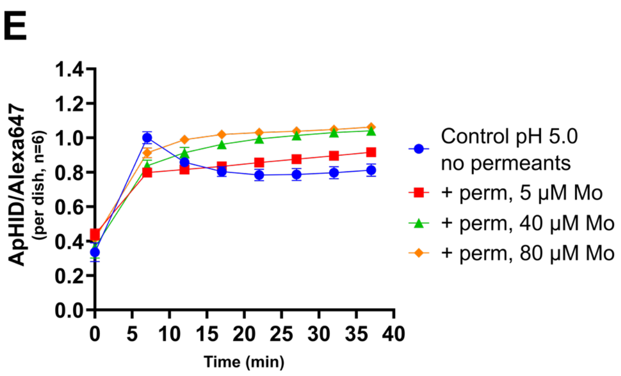


**Suppl. Fig S2. Supplementation of TRIS maleate pH 5.0 buffer with monensin facilitates buffer equilibration across membranes at 37 ºC**. **(A-D)** Fixed J774 macrophage lysosomes loaded with ApHID-Alexa 647 dextrans imaged by confocal microscopy. Cells were imaged in PBS (time 0, A-D) and then in 50 mM TRIS maleate pH 5.0 buffer alone (A) or supplemented with 40 mM methylamine, 40 mM sodium acetate and 5 µM (B), 40 µM (C), or 80 µM (D) monensin. Cells were imaged every 5 min for 40 min. Overlays show ApHID (green) and Alexa 647 (red) channels for each buffer condition. **(E)** ApHID/Alexa 647 ratios measured for each buffer condition normalized to 80 µM condition (36 min time point), plotted against incubation time. Two wells were imaged per condition, 3 fields acquired per well. The experiment was repeated three times. Geometrical objects and bars indicate averages ±SEM. Scale bars: 10 µm. Abbreviations: 'perm': membrane-permeant equilibrators. 'A647': Alexa Fluor 647.

Suppl. Fig. S2 shows fluorescence measurements of ApHID (green) and Alexa 647 (red) in macrophages incubated with buffers. Buffers supplemented with various amounts of monensin also contained 40 mM methylamine and 40 mM sodium acetate as membrane-permeant equilibrators.

Monensin supplementation led to increased ApHID fluorescence relative to cells in buffer without membrane permeants (2SB-2SD vs 2SA). 40 μM and 80 μM monensin addition led to ApHID/Alexa647 stabilization within 30 mins of incubation, whereas absence of membrane permeants altogether led to incomplete ratio equilibration (S2E). Based on that, the supplementation of 50 mM TRIS maleate pH 5.0 buffer with 40-80 μM monensin, together with methylamine and sodium acetate, was found to be effective at ensuring optimal buffer diffusion across membranes in fixed cells at 37 ºC.

#### Ratio-to-pH calibration curves for ApHID, fluorescein and Oregon Green (supporting main section Figure 6 and 7).

Suppl. Fig. 3 shows averaged ratio-to-pH calibrations for the ApHID/Alexa647, fluorescein/Alexa647, and OG/Alexa647 pairs, obtained by measuring dextrans in solution using a SpectraMax M3 spectrophotometer as described in the Methods section. These calibrations result from averaging 2 individual titrations for each dye pair. Titration data in solution is used to prepare full calibration curves for fluorescence-to-pH interpolation (Figures 6 and 7 of the main text). Titrations were fit to a 4-component sigmoidal curve (Suppl. Fig 3A). Alexa 647 was generally pH-independent in the pH range of 4.5-6.5 (Suppl. Fig. 3B).

To prepare a ratio-to-pH calibration for ratiometric pH imaging, J774 macrophages seeded on 96-well plates were fixed in 0.5% PFA and incubated in 50 mM TRIS maleate pH 5.0 buffer containing 10% FBS, 40 mM methylamine hydrochloride, 40 mM sodium acetate, and 40 μM monensin for 30 min at 37 °C, followed by confocal microscopy imaging. Measured pH-sensitive dye/Alexa 647 ratios corresponding to pH 5.0 in fixed cells were thereafter used to generate all subsequent ratios for pH values 3.5 and 5.0-7.4, using the titration data corresponding to Suppl. Fig. S3A.

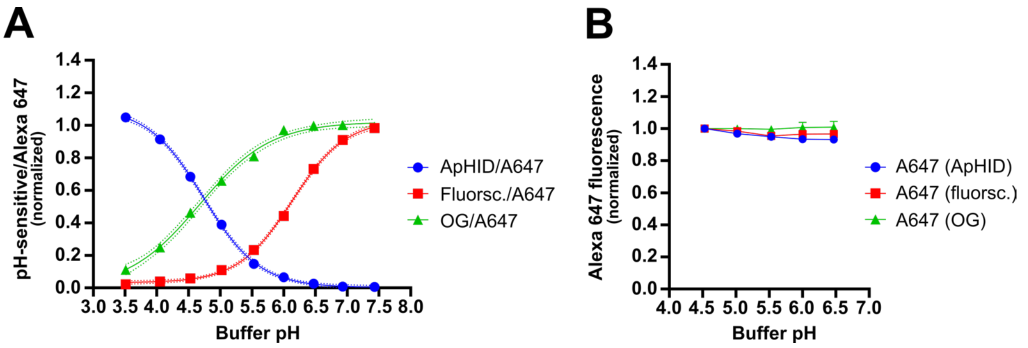


**Supplementary Figure 3. Average pH-sensitive/Alexa 647 vs. pH titrations used to prepare ratio-to-pH calibrations for ratiometric pH imaging experiments on J774 macrophages. (A)** pH-sensitive/pH-independent vs. pH titrations for dextrans labeled with one of the pH-sensitive probes as well as Alexa 647 (pH-independent). To prepare each individual curve, dextrans were solubilized in buffers with pH adjusted between 3.5 and 7.5 containing 10% FBS, 40 mM methylamine hydrochloride, 40 mM sodium acetate and 40 uM monensin, and measured at 37°C using a SpectraMax M3 spectrophotometer (Molecular Devices). The resulting ratio-to-pH titrations were fit to 4-component sigmoidal curves (superimposed colored curves). **(B)** Alexa 647 fluorescence plotted against buffer pH for each dextran is generally pH-independent in the pH range of 4.5-6.5. Dextrans were solubilized in buffers, loaded into 384-well plates in triplicate, and measured using a SpectraMax M3 plate reader (see Methods section). See Suppl. Table ST3 for statistics on calibration. Measurements in solution were repeated twice. Geometric figures and bars represent averaged ratios ±SEM. The SEM error bars fit within the symbols. Superimposed black dotted lines represent standard error of the sigmoidal fit and are superimposed on the curves.

| Supplementary Table ST1. Photostability of ApHID, fluorescein and Oregon Green conjugated to 70 KDa dextrans. Statistics. | | | | | | | | | |
|---|---|---|---|---|---|---|---|---|---|
| | ApHID (70KDa dextran) | | | Fluorescein (70KDa dextran) | | | OG (70KDa dextran) | | |
| Irradiation cycle | Norm. $F_0/F$ | SD | n (fields) | Norm. $F_0/F$ | SD | n (fields) | Norm. $F_0/F$ | SD | n (fields) |
| 1 | 1.00 | 0.00 | 16 | 1.00 | 0.00 | 16 | 1.00 | 0.00 | 16 |
| 2 | 1.00 | 0.00 | 16 | 0.90 | 0.02 | 16 | 0.91 | 0.01 | 16 |
| 3 | 1.00 | 0.00 | 16 | 0.83 | 0.03 | 16 | 0.83 | 0.02 | 16 |
| 4 | 0.99 | 0.01 | 16 | 0.76 | 0.04 | 16 | 0.77 | 0.03 | 16 |
| 5 | 0.99 | 0.01 | 16 | 0.71 | 0.04 | 16 | 0.71 | 0.03 | 16 |
| 6 | 0.99 | 0.01 | 16 | 0.66 | 0.05 | 16 | 0.67 | 0.03 | 16 |
| 7 | 0.99 | 0.01 | 16 | 0.62 | 0.05 | 16 | 0.63 | 0.04 | 16 |
| 8 | 0.98 | 0.01 | 16 | 0.59 | 0.05 | 16 | 0.60 | 0.04 | 16 |
| 9 | 0.98 | 0.01 | 16 | 0.56 | 0.05 | 16 | 0.56 | 0.04 | 16 |
| 10 | 0.98 | 0.01 | 16 | 0.53 | 0.05 | 16 | 0.54 | 0.04 | 16 |
| 11 | 0.98 | 0.01 | 16 | 0.50 | 0.05 | 16 | 0.51 | 0.04 | 16 |
| 12 | 0.97 | 0.01 | 16 | 0.48 | 0.05 | 16 | 0.49 | 0.04 | 16 |
| 13 | 0.97 | 0.01 | 16 | 0.46 | 0.05 | 16 | 0.47 | 0.04 | 16 |
| 14 | 0.97 | 0.02 | 16 | 0.44 | 0.05 | 16 | 0.45 | 0.04 | 16 |
| 15 | 0.97 | 0.02 | 16 | 0.42 | 0.05 | 16 | 0.43 | 0.04 | 16 |
| 16 | 0.96 | 0.02 | 16 | 0.41 | 0.05 | 16 | 0.42 | 0.04 | 16 |
| 17 | 0.96 | 0.02 | 16 | 0.39 | 0.04 | 16 | 0.40 | 0.04 | 16 |
| 18 | 0.96 | 0.02 | 16 | 0.38 | 0.04 | 16 | 0.39 | 0.04 | 16 |
| 19 | 0.96 | 0.02 | 16 | 0.36 | 0.04 | 16 | 0.37 | 0.04 | 16 |
| 20 | 0.95 | 0.02 | 16 | 0.35 | 0.04 | 16 | 0.36 | 0.04 | 16 |
| 21 | 0.95 | 0.02 | 16 | 0.34 | 0.04 | 16 | 0.35 | 0.04 | 16 |
| 22 | 0.95 | 0.02 | 16 | 0.33 | 0.04 | 16 | 0.34 | 0.04 | 16 |
| 23 | 0.94 | 0.02 | 16 | 0.32 | 0.04 | 16 | 0.33 | 0.04 | 16 |
| 24 | 0.94 | 0.02 | 16 | 0.31 | 0.04 | 16 | 0.32 | 0.04 | 16 |
| 25 | 0.94 | 0.02 | 16 | 0.30 | 0.04 | 16 | 0.31 | 0.04 | 16 |
| 26 | 0.94 | 0.02 | 16 | 0.29 | 0.04 | 16 | 0.30 | 0.04 | 16 |
| 27 | 0.93 | 0.02 | 16 | 0.28 | 0.04 | 16 | 0.29 | 0.04 | 16 |
| 28 | 0.93 | 0.02 | 16 | 0.27 | 0.04 | 16 | 0.29 | 0.04 | 16 |
| 29 | 0.93 | 0.02 | 16 | 0.27 | 0.04 | 16 | 0.28 | 0.04 | 16 |
| 30 | 0.92 | 0.02 | 16 | 0.26 | 0.03 | 16 | 0.27 | 0.03 | 16 |
| 31 | 0.92 | 0.02 | 16 | 0.25 | 0.03 | 16 | 0.26 | 0.03 | 16 |
| 32 | 0.92 | 0.02 | 16 | 0.25 | 0.03 | 16 | 0.26 | 0.03 | 16 |
| 33 | 0.92 | 0.02 | 16 | 0.24 | 0.03 | 16 | 0.25 | 0.03 | 16 |
| 34 | 0.91 | 0.02 | 16 | 0.24 | 0.03 | 16 | 0.25 | 0.03 | 16 |
| 35 | 0.91 | 0.02 | 16 | 0.23 | 0.03 | 16 | 0.24 | 0.03 | 16 |
| 36 | 0.91 | 0.03 | 16 | 0.22 | 0.03 | 16 | 0.24 | 0.03 | 16 |
| 37 | 0.91 | 0.03 | 16 | 0.22 | 0.03 | 16 | 0.23 | 0.03 | 16 |
| 38 | 0.90 | 0.03 | 16 | 0.21 | 0.03 | 16 | 0.23 | 0.03 | 16 |
| 39 | 0.90 | 0.03 | 16 | 0.21 | 0.03 | 16 | 0.22 | 0.03 | 16 |
| 40 | 0.90 | 0.03 | 16 | 0.21 | 0.03 | 16 | 0.22 | 0.03 | 16 |
| 41 | 0.90 | 0.03 | 16 | 0.20 | 0.03 | 16 | 0.21 | 0.03 | 16 |
| 42 | 0.89 | 0.03 | 16 | 0.20 | 0.03 | 16 | 0.21 | 0.03 | 16 |
| 43 | 0.89 | 0.03 | 16 | 0.19 | 0.03 | 16 | 0.20 | 0.03 | 16 |
| 44 | 0.89 | 0.03 | 16 | 0.19 | 0.03 | 16 | 0.20 | 0.03 | 16 |
| 45 | 0.89 | 0.03 | 16 | 0.19 | 0.03 | 16 | 0.20 | 0.03 | 16 |
| 46 | 0.89 | 0.03 | 16 | 0.18 | 0.03 | 16 | 0.19 | 0.03 | 16 |
| 47 | 0.88 | 0.03 | 16 | 0.18 | 0.03 | 16 | 0.19 | 0.03 | 16 |
| 48 | 0.88 | 0.03 | 16 | 0.17 | 0.03 | 16 | 0.19 | 0.03 | 16 |
| 49 | 0.88 | 0.03 | 16 | 0.17 | 0.03 | 16 | 0.18 | 0.03 | 16 |
| 50 | 0.88 | 0.03 | 16 | 0.17 | 0.03 | 16 | 0.18 | 0.03 | 16 |

**Supplementary Section Table ST1 supporting main section Fig. 3 .** Lysosomes in fixed J774 macrophages, loaded with 10 KDa dextrans labeled with ApHID, fluorescein or Oregon Green, were exposed to 488 nm laser irradiation for 50 half-second pulses. Green fluorescence intensity was measured at time 0 ($F_0$) and after each irradiation cycle, separated by 1 second intervals (F). Fluorescence was normalized to time 0 ($F_0$) measurement. The experiment was repeated twice; two dishes were measured per condition and 4 fields were imaged per dish for a total of 16 fields. Average $F/F_0$ fluorescence ±SD is presented. Abbreviations: 'OG': Oregon Green.

**Supplementary Table ST2 supporting Figure 4. Descriptive statistics**

| ApHID:Alexa 405 | ApHID pKa | ApHID:Cy5•3SO3⁻ | ApHID pKa | Charge (ApHID:Alexa 405) | ApHID pKa |
|---|---|---|---|---|---|
| **1 : 2** | 5.08 | **1.2 : 1.4** | 5.10 | **Control no Alexa405** | 5.07 |
| **1 : 1** | 5.04 | **2.3 : 1.4** | 5.03 | **1 : 1.33** | 5.00 |
| **3 : 2** | 5.06 | **3.2 : 1.4** | 5.04 | **1 : 6** | 5.12 |
| **2 : 1** | 5.00 | **4.6 : 1.4** | 5.07 | **1 : 22** | 5.42 |

**Supplementary Section Table ST2 supporting main section Fig. 4.** Effect of various amounts of pH-independent fluorophores and dextran polymer charge density on ApHID pKa. Dextrans were labeled with ApHID and varying degrees of Cy5•3SO3⁻ and Alexa405. The amounts of Cy5•3SO3⁻ and Alexa405 per molecule of dextran, relative to ApHID, are presented together with their corresponding ApHID pKa measured for that particular dextran. Charge density was introduced by labeling the polymers with various amounts of Alexa 405, which carries three negatively charged sulfate groups. Experiments were repeated three times. Average ApHID pKa values are presented.

**Supplementary Table ST3 supporting Supplementary Figure 3. Descriptive Statistics**

| | ApHID-A647 dex +FBS @37C | | | Fluorsc-A647 dex +FBS @37C | | | OG-A647 dex +FBS @37C | | |
|---|---|---|---|---|---|---|---|---|---|
| **pH buffer** | **Norm. ApHID/A647** | **SEM** | **n** | **Norm. fluorsc./A647** | **SEM** | **n** | **Norm. OG/A647** | **SEM** | **n** |
| **3.51** | 1.05 | 0.02 | 2 | 0.02 | 0.00 | 2 | 0.11 | 0.00 | 2 |
| **4.05** | 0.91 | 0.01 | 2 | 0.04 | 0.00 | 2 | 0.25 | 0.00 | 2 |
| **4.53** | 0.68 | 0.01 | 2 | 0.06 | 0.00 | 2 | 0.46 | 0.01 | 2 |
| **5.02** | 0.39 | 0.00 | 2 | 0.11 | 0.01 | 2 | 0.66 | 0.00 | 2 |
| **5.53** | 0.15 | 0.01 | 2 | 0.23 | 0.01 | 2 | 0.81 | 0.01 | 2 |
| **6** | 0.06 | 0.00 | 2 | 0.44 | 0.00 | 2 | 0.97 | 0.01 | 2 |
| **6.47** | 0.03 | 0.00 | 2 | 0.73 | 0.00 | 2 | 1.00 | 0.01 | 2 |
| **6.93** | 0.01 | 0.00 | 2 | 0.91 | 0.01 | 2 | 1.00 | 0.00 | 2 |
| **7.43** | 0.01 | 0.00 | 2 | 0.98 | 0.01 | 2 | 0.99 | 0.00 | 2 |

| | ApHID-A647 dex +FBS @37C | | | Fluorsc-A647 dex +FBS @37C | | | OG-A647 dex +FBS @37C | | |
|---|---|---|---|---|---|---|---|---|---|
| **pH buffer** | **Norm. A647 fluo.** | **SEM** | **n** | **Norm. A647 fluo.** | **SEM** | **n** | **Norm. A647 fluo.** | **SEM** | **n** |
| **3.51** | 1.05 | 0.00 | 2 | 1.06 | 0.03 | 2 | 1.02 | 0.01 | 2 |
| **4.05** | 1.03 | 0.02 | 2 | 1.02 | 0.02 | 2 | 1.02 | 0.01 | 2 |
| **4.53** | 1.00 | 0.00 | 2 | 1.00 | 0.00 | 2 | 1.00 | 0.00 | 2 |
| **5.02** | 0.97 | 0.00 | 2 | 0.98 | 0.02 | 2 | 1.00 | 0.01 | 2 |
| **5.53** | 0.95 | 0.02 | 2 | 0.95 | 0.02 | 2 | 1.00 | 0.02 | 2 |
| **6** | 0.93 | 0.00 | 2 | 0.97 | 0.02 | 2 | 1.01 | 0.03 | 2 |
| **6.47** | 0.93 | 0.01 | 2 | 0.97 | 0.02 | 2 | 1.01 | 0.04 | 2 |
| **6.93** | 0.92 | 0.00 | 2 | 0.96 | 0.01 | 2 | 0.97 | 0.01 | 2 |
| **7.43** | 0.91 | 0.00 | 2 | 0.99 | 0.01 | 2 | 0.98 | 0.02 | 2 |

**Supplementary Table ST3 supporting Supplementary Figure S3.** Normalized average pH-sensitive/Alexa647 ratios and Alexa647 fluorescence intensity vs. pH titrations measured for dextrans in solution, plotted in Supplementary Figure S3. Averaged ratio titrations for each type of dextran (from two measurements in solution) were used to prepare ratio-to-pH calibrations for ratiometric pH imaging experiments on J774 macrophages (main section Fig. 6 and Fig. 7). Average Alexa647 intensity is generally pH-independent. Abbreviations: fluorsc, fluorescein; OG, Oregon Green; A647, Alexa 647.

| Supplementary Table ST4 supporting main section Figure 6. Statistics | | | | | | |
|---|---|---|---|---|---|---|
| | ApHID | ApHID +MeNH$_2$ | Fluorescein | Fluorescein +MeNH$_2$ | Oregon Green | OG +MeNH$_2$ |
| **Average pH ±SEM** | 4.99 ± 0.03 | 6.42 ± 0.01 | 5.06 ± 0.01 | 6.28 ± 0.01 | 4.98 ± 0.01 | 6.56 ± 0.13 |

**Unpaired One-way ANOVA (Dunnett's multiple comparison test)**

| *Comparison* | *Mean 1* | *Mean 2* | *Mean Diff.* | *p value* | *Significance* |
|---|---|---|---|---|---|
| ApHID vs. Fluorescein | 4.99 | 5.06 | -0.07 | 0.3 | n.s. |
| ApHID vs. OG | 4.99 | 4.98 | 0.02 | 0.97 | n.s. |
| ApHID vs ApHID +MeNH2 | 4.99 | 6.42 | -1.43 | <0.0001 | **** |
| Fluorescein vs. Fluoc. +MeNH2 | 5.06 | 6.28 | -1.21 | <0.0001 | **** |
| OG vs. OG. +MeNH2 | 4.98 | 6.56 | -1.6 | <0.001 | *** |

**Supplementary Section Table ST4 supporting main section Fig. 6 .** Lysosomal pH ± SEM reported by ratiometric pH imaging of J774 lysosomes using ApHID, fluorescein and Oregon Green dextrans, also labeled with Alexa 647 (pH-independent) with statistical comparisons. Average pH values were calculated using a total of six wells imaged in three independent experiments. Two wells were imaged for each condition, and four fields were imaged per dish. Differences in averaged pH between probes and conditions were assessed using the unpaired One-way ANOVA with Dunnett’s multiple comparison test using 95% confidence interval. P-values shown as $p>0.05$ (ns), $p\leq0.05$ (*), $p\leq0.01$ (**), $p\leq0.001$ (***), and $p\leq0.0001$ (****). Abbreviations: ‘OG’: Oregon Green; ‘Fluoc’: fluorescein.

**Supplementary Table ST5 supporting main section Figure 7. Descriptive statistics.**

| | ApHID | | | Fluorescein | | | OG | | |
|---|---|---|---|---|---|---|---|---|---|
| **Time (min)** | **LE/Ly pH** | **SEM** | **n** | **LE/Ly pH** | **SEM** | **n** | **LE/Ly pH** | **SEM** | **n** |
| **20** | 5.03 | 0.07 | 6 | 5.05 | 0.06 | 6 | 4.93 | 0.11 | 6 |
| **25** | 5.02 | 0.10 | 6 | 5.07 | 0.04 | 6 | 4.98 | 0.06 | 6 |
| **30** | 5.01 | 0.11 | 6 | 5.07 | 0.03 | 6 | 4.99 | 0.03 | 6 |
| **35** | 4.99 | 0.10 | 6 | 5.07 | 0.03 | 6 | 4.99 | 0.02 | 6 |
| **40** | 4.99 | 0.09 | 6 | 5.07 | 0.03 | 6 | 4.99 | 0.02 | 6 |
| **45** | 4.99 | 0.09 | 6 | 5.07 | 0.03 | 6 | 4.99 | 0.02 | 6 |
| **50** | 4.99 | 0.08 | 6 | 5.07 | 0.03 | 6 | 4.98 | 0.02 | 6 |
| **55** | 4.99 | 0.08 | 6 | 5.06 | 0.03 | 6 | 4.98 | 0.02 | 6 |
| **60** | 4.99 | 0.08 | 6 | 5.06 | 0.03 | 6 | 4.98 | 0.02 | 6 |
| **65** | 4.99 | 0.08 | 6 | 5.06 | 0.02 | 6 | 4.98 | 0.02 | 6 |
| **70** | 4.99 | 0.08 | 6 | 5.06 | 0.02 | 6 | 4.98 | 0.02 | 6 |
| **75** | 4.99 | 0.08 | 6 | 5.06 | 0.03 | 6 | 4.97 | 0.02 | 6 |

**Supplementary Section Table ST5 supporting main section Fig. 7 .** Real-time ratiometric imaging of J774 macrophage lysosomal pH reported over the course of a continuous 75 min acquisition. Macrophage lysosomes were loaded with dextrans labeled with ApHID, fluorescein or Oregon Green (pH-sensitive) and Alexa 647 (pH-independent). Images were acquired every 5 min for a total imaging time of 75 min. Average pH values were calculated using a total of six wells imaged in three independent experiments. Two wells were imaged for each condition, and four fields were imaged per dish. Average pH ±SEM is presented. Abbreviations: ‘OG’: Oregon Green.

| Dextran weight (KDa) / Labeling conc. mg/mL | *NHS ester excess molar ratio (reacted vs. incorporated)* | | | | | | | |
|---|---|---|---|---|---|---|---|---|
| | **Vendor** | **Experiment** | **x ApHID** | **x fluorescein** | **x Oregon Green** | **x Alexa 405** | **x Cy5-$3xSO_3$** | **x Alexa 647** |
| 10 / 50 | TS | *General spectroscopy* | 2.35 (1.6) | 1.43 (1.2) | 1.43 (1.2) | - | - | |
| 70 / 20 | Fina | *Derivatization* | 2 (1), 4 (2), 6 (3), 8 (4) | - | - | 4 (2) | 4 (1.4) | |
| 70 / 20 | Fina | *Net charge effect* | 3 (1.5) | - | - | 3 (1.3), 15 (6), 100 (22) | - | |
| 70 / 25 | TS | *Photobleaching* | 3 (2.1) | 3 (1.8) | 3 (1.9) | - | - | |
| 70 / 25 | TS | *Ratiometric pH imaging* | 3 (1.6) | 3 (1.8) | - | 3 (1.8) | - | |
| | | | 4 (2.6) | 4 (1.69) | 4 (2.06) | - | 4-6 (2.8-3) | 3 (1.8-2.2) |

**Supplementary Section Table ST6. List of dextrans used for each experiment in the study.** Information for each dextran used in our experiments is shown, including dextran weight (KDa), concentration during fluorophore labeling (mg/mL), vendor, type of fluorophore attached and dextran:NHS probe molar ratio used for labeling (black characters), with their resulting labeling after dialysis (red characters). Abbreviations: TS (Thermo Fisher); Fina (Fina Biosolutions), KDa (kiloDalton); mg/mL (milligram/milliliter).

#### Key Resource Table

| Resource | Product | Source | Identifier |
|---|---|---|---|
| **Chemical** | | | |
| Buffer salt | Sodium hydroxide | Sigma-Aldrich | S5881 |
| Buffer salt | Sodium phosphate monobasic anhydrous | Sigma-Aldrich | S8262 |
| Buffer salt | Calcium chloride dihydrate | Sigma-Aldrich | C5080 |
| Buffer salt | Magnesium chloride hexahydrate | Sigma-Aldrich | 102510804 |
| Buffer salt | Sodium acetate anhydrous | Sigma-Aldrich | 58750 |
| Buffer salt | Sodium phosphate dibasic heptahydrate | Sigma-Aldrich | 59390 |
| Buffer salt | Citric acid anhydrous | Sigma-Aldrich | C4540 |
| Buffer salt | Sodium citrate trisodium salt dihydrate | Sigma-Aldrich | S4641 |
| Buffer salt | Trizma Maleate | Sigma-Aldrich | T3128 |
| Buffer salt | Trizma Base | Sigma-Aldrich | T1503 |
| Buffer salt | Trizma Hydrochloride | Roche | 10812846001 |
| Buffer salt | Sodium bicarbonate | Sigma-Aldrich | S5761 |
| Buffer salt | HEPES | Sigma-Aldrich | H3375 |
| Buffer salt | Sodium chloride | Sigma-Aldrich | S9625 |
| Buffer salt | Potassium chloride | Sigma-Aldrich | P5405 |
| Reagent | Iron (II) perchlorate hydrate | Sigma-Aldrich | 334081 |
| Reagent | Hydrogen peroxide 30% solution | VWR | BDH7690-3 |
| Reagent | Bovine serum albumin | Sigma-Aldrich | A2153 |
| Reagent | Paraformaldehyde 32% solution | Electron Microscopy Sciences | 15714-S |
| Perm. Equilibrator | Monensin Sodium Salt | Sigma-Aldrich | M5273 |
| Perm. Equilibrator | Nigericin Sodium Salt | Sigma-Aldrich | N7143 |
| Perm. Equilibrator | Methylamine Hydrochloride | Sigma-Aldrich | M0505 |
| **Fluorophores** | | | |
| Fluorophore | NHS Alexa 405 | Thermo Fisher | A30000 |
| Fluorophore | NHS Alexa 647 | Thermo Fisher | A20106 |
| Fluorophore | NHS Cy5•3x$SO_3^-$ (Cy5 SE TRI SO3) | AstaTech | 44193 |
| Fluorophore | NHS Acidic pH Indicator Dye (ApHID) | Custom-made | - |
| Fluorophore | NHS 5/6-carboxyfluorescein | Thermo Fisher | 46410 |
| Fluorophore | NHS Oregon Green | Thermo Fisher | O6147 |
| Fluorophore | LysoSensor yellow/blue 10 KDa | Thermo Fisher | L22460 |
| Dextran | Amino Dextran polymer 10 KDa | Thermo Fisher | D1860 |
| Dextran | Amino Dextran polymer 70 KDa | Thermo Fisher | D1862 |
| Dextran | Amino Dextran polymer 70 KDa | Fina Biosolutions | AD70x33 |
| **Cells** | | | |
| Cell line | Murine macrophages (sarcoma) | ATCC | J774.A1 TIB-67 |
| **Growth Media** | | | |
| Growth medium | Dubbelco's Modified Eagle Medium | Corning | 15-013-CV |
| Growth medium | Dubbelco's Modified Eagle Medium (no phenol red) | Corning | 90-013-PB |

| | | | |
|---|---|---|---|
| Growth factor | Fetal Bovine Serum (FBS) | Gemini | 100-106 |
| Growth factor | L-glutamine | Gibco | 25030081 |
| Growth factor | D-(+)-glucose | Sigma-Aldrich | G7021 |
| Growth factor | Sodium pyruvate | Sigma-Aldrich | S8636 |
| Antibiotic | Penicillin-Streptomycin | Thermo Fisher | 15140163 |
| **Lab equipment** | | | |
| Purification | 3.5 KDa Side-A-Lyzer dialysis cassettes | Thermo Fisher | 66330 |
| Purification | 20 KDa Side-A-Lyzer dialysis cassettes | Thermo Fisher | 66003 |
| Purification | Ultra-pure distilled water | Hydro Services | PicoPure3 System |
| Purification | Filtration cups, 0.2 µm aPES (0.5 L) | Thermo Fisher | 595-3320 |
| Purification | Filtration cups, 0.2 µm aPES (1 L) | Thermo Fisher | 597-4520 |
| Spectrophotometer | Spectrophotometer / plate reader | Molecular Devices | SpectraMax M3 |
| Spectrophotometer | 384-well polystyrene microplates | Corning | 3746 |
| Incubation system | Gravity convection oven (Isotemp) | Fisher Scientific | 15-103-0503 |
| Confocal | Confocal microscope | Zeiss | LSM 880 |
| Confocal | Confocal microscope | Leica | Stellaris |
| Optics | Power Meter | Coherent | LaserMate Q |
| **Computer software** | | | |
| Data processing | Statistical data processor | Microsoft | Excel 365 v. 2208 |
| Data processing | Statistical data processor | GraphPad Software | GraphPad Prism v. 9.0 |
| Data processing | Digital image processor | Molecular Devices | MetaMorph v. 6.7.1.157 |
| Data processing | Word processor | Microsoft | MS Word 365 v. 2208 |
| Data processing | Image editor | Inkscape | Inkscape v. 1.2.2 |
| Data processing | Chemical drawing | PerkinElmer | Inkscape v. 1.2.2 |

### ApHID Synthetic Protocol

#### 1. Complete list of reagents and equipment used in ApHID synthetic procedures.

| Resource | Product | Source | Identifier |
|---|---|---|---|
| **Chemical** | | | |
| Reagent | *N*-[(Dimethylamino)-1*H*-1,2,3-triazolo-[4,5-*b*]pyridin-1-ylmethylene]-*N*-methylmethanaminium hexafluorophosphate *N*-oxide (HATU) | Sigma | 445460 |
| Reagent | Ethylamine (2M in THF) | Sigma | 395072 |
| Reagent | $NH_2$-$PEG_4$-COOH | Sigma | QBD10244 |
| Reagent | EDC | Sigma | 39391 |
| Reagent | *N*-hydroxysuccinimide | Sigma | 130672 |
| Drying agent | Sodium sulfate, anhydrous | Sigma | 239313 |
| Solvent | Dimethylformamide | Sigma | 227056 |
| Solvent | Methanol | Sigma | 439193 |
| Solvent | Triethylamine | Sigma | 471283 |
| Solvent | Dichloromethane | Sigma | 650463 |
| Solvent | Dimethylsulfoxide-$d_6$ | Sigma | 151874 |
| Solvent | Tetrahydrofuran | Sigma | 401757 |
| Solvent | Acetonitrile | Sigma | AX0156 |
| Solvent | Formic Acid | Sigma | 5330020050 |
| **Lab equipment** | | | |
| Purification | CombiFlash Rf+ | Teledyne ISCO | 66330 |
| Purification | Silica gel column | Teledyne ISCO | 69-2203-344 |
| Purification | C18 column | Teledyne ISCO | 69-2203-328 |
| Drying | Lyophylizer | Labconco | 7934027 |
| NMR | Avance III HD 500 MHz | Bruker | |
| **Computer software** | | | |
| Graphics | Chemical drawing | PerkinElmer | ChemDraw v 22.2.0 |
| Data analysis | NMR data analysis | Mestrelab Research | Mnova v 14.3.2 |

**Supplementary Table 3. List of reagents and equipment used in ApHID's synthetic procedures.**

#### 2. ApHID Chemical Synthesis protocol

**Synthesis of known precursors**

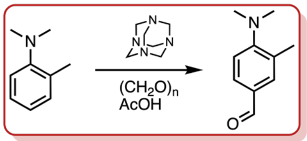


**Reference:** Gawinecki et al. [3]

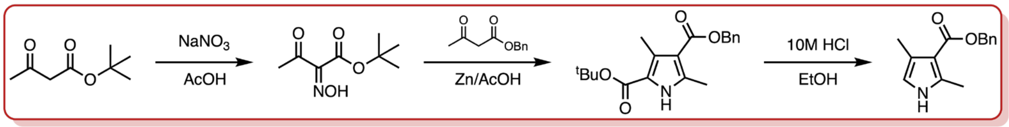


**Reference:** Li et al. [4]

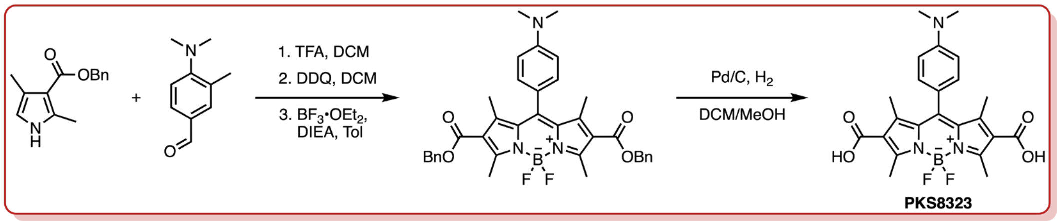


**Reference:** Maeda et al. [5]

**Synthesis of new molecules.**

**General Information:** Unless otherwise stated, all commercially available materials were purchased from Sigma Aldrich and were used without further purification. When necessary, solvents and reagents were dried prior to use, using standard protocols. All non-aqueous reactions were carried out in oven-dried glassware under an atmosphere of Argon. $^{1}H$ and $^{13}C$ NMR spectra were acquired on a Bruker Avance III HD spectrometer at 500 MHz for $^{1}H$ and 125 MHz for $^{13}C$. Chemical shifts are expressed in parts per million downfield from tetramethylsilane (TMS), using either TMS or the solvent resonance as an internal standard (TMS, $^{1}H$: 0 ppm; chloroform, $^{13}C$: 77.0 ppm; DMSO-$d_6$, $^{1}H$: 2.5 ppm; $^{13}C$: 39.5 ppm). Data are reported as follows: chemical shift, multiplicity (s = singlet, d = doublet, t = triplet, q = quartet, m = multiplet, br = broad), integration, and coupling constant. LC-MS analysis using a Waters I-Class ACQUITY UPLC system equipped with an ACQUITY Photodiode Array (PDA), a Waters SQD2 mass spectrometer, and a Waters ACQUITY BEH C18 column (1.7 µm, 2.1 × 100 mm). The solvent system consisted of 0.1% formic acid in water (solvent A) and 0.1% formic acid in acetonitrile (solvent B). Flow was set to 0.3 ml/min, and a gradient of 5% to 95% solvent B was applied over a period of 3 minutes. The total run time was 4 minutes. Eluents were detected using a PDA at a wavelength of 254 nm. Mass data were obtained in both positive and negative electrospray mode at a cone voltage of 30 V. HPLC purifications were performed using a Waters AutoPure HPLC/MS system equipped with XBridge OBD prep C18 5µm (19 x 150 mm) column and SQD2 mass spectrometer.

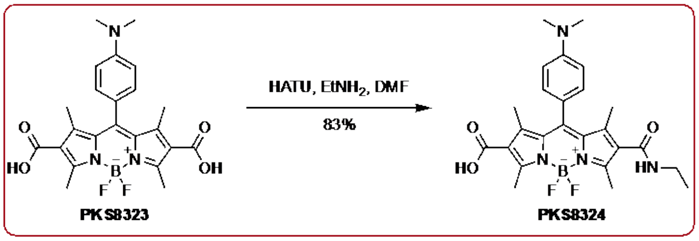


#### Synthesis of PKS8324

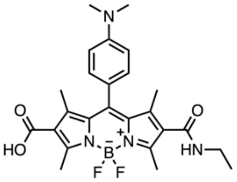

PKS8323 (2.00 g, 2.98 mmol, 2•TEA salt) was dissolved in DMF (20 mL), and the solution was cooled to 0 °C. *N*-[(dimethylamino)-1*H*-1,2,3-triazolo-[4,5-*b*]pyridin-1-ylmethylene]-*N*-methylmethanaminium hexafluorophosphate *N*-oxide (HATU) (1.13 g, 2.98 mmol) was added to the solution at 0 °C. After stirring for 5 minutes at 0 °C, ethylamine (2M in THF, 9.0 mmol, 4.5 mL) was added, and the mixture was allowed to warm to ambient temperature slowly. The reaction gave a mixture of starting material, desired product, and diamide. The mixture was purified by CombiFlash [silica gel; 0-10% methanol (0.5% $Et_3N$) in DCM (0.5% $Et_3N$)] to give the product (5 eq. triethylamine by NMR; 2.47 g, 83%) as a red solid. $^1$H NMR (500 MHz, DMSO-$d_6$) δ 1.05 (t, *J* = 7.2 Hz, 3H), 1.43 (s, 3H), 1.66 (s, 3H), 2.31 (s, 3H), 2.51 (s, 3H), 2.69 (s 3H), 2.71 (s, 6H), 3.14 – 3.22 (m, 2H), 7.08 – 7.11 (m, 2H), 7.17 (d, *J* = 8.0 Hz, 1H), 8.12 (t, *J* = 5.5 Hz, 1H), 10.21 (br, 1H).

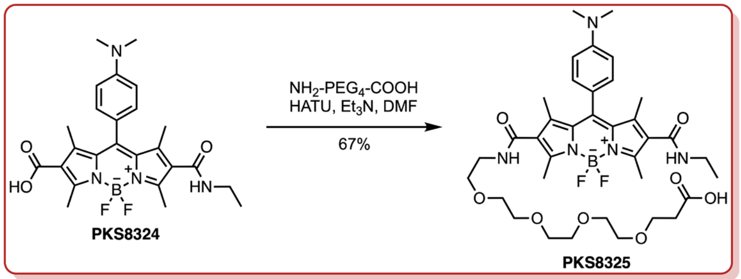


#### Synthesis of PKS8325

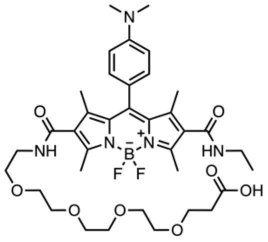

PKS8324 (2.47 g, 2.47 mmol, 5 eq TEA) and HATU (971 mg, 2.55 mmol) were dissolved in DMF (15 mL) under an argon atmosphere. The solution was cooled to 0 °C, and $NH_2$-PEG$_4$-COOH (645 mg, 2.43 mmol) was added. After stirring for 5 minutes at 0 °C, triethylamine (4.87 mmol, 680 μL) was added. The reaction mixture was allowed to warm to ambient temperature and stirred overnight. After completion of the reaction, the solvent was evaporated, and the mixture was purified by CombiFlash [silica gel; 0–30% methanol (0.5% $Et_3N$) in DCM (0.5% $Et_3N$)] to give the product (1.21 g, 67%) as a red-brown solid. $^1$H NMR (500 MHz, DMSO-$d_6$) δ 1.05 (t, *J* = 7.2 Hz, 3H), 1.43 (s, 3H), 1.44 (s, 3H), 2.32 (s, 3H), 2.34 (t, *J* = 6.6 Hz, 2H), 2.50 ( 6H), 2.71 (s, 6H), 3.15 – 3.21 (m, 2H), 3.25 – 3.50 (m, 16H), 3.56 (t, *J* = 6.6 Hz, 2H), 7.06 – 7.08 (m, 2H), 7.18 (d, *J* = 8.1 Hz, 1H), 8.11 (t, *J* = 6.0 Hz, 1H), 8.13 (t, *J* = 5.3 Hz, 1H).

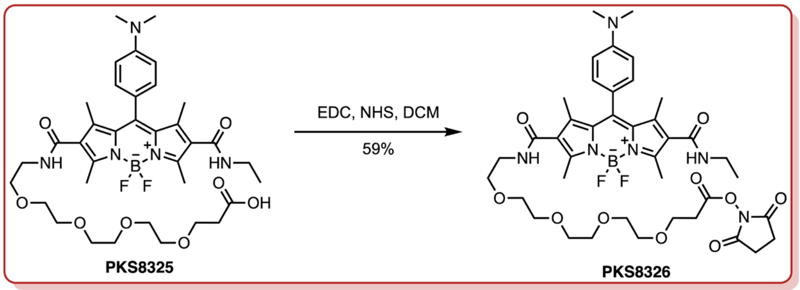


**Synthesis of PKS8326**

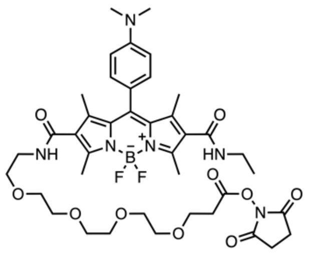

PKS8325 (1.21 g, 1.65 mmol) and EDC (575 mg, 3.00 mmol) were dissolved in DCM (50 mL) under an argon atmosphere. The solution was cooled to 0 °C, and *N*-hydroxysuccinimide (259 mg, 2.25 mmol) was added. The reaction mixture was allowed to warm to ambient temperature slowly and stirred at ambient temperature overnight. The reaction mixture was diluted with water and extracted with dichloromethane. The organic layer was washed with saturated brine solution, dried over anhydrous $Na_2SO_4$, and evaporated. The crude was purified by CombiFlash (C-18 column; 0% – 100% acetonitrile in water). The fractions with the product were combined, the acetonitrile was evaporated, and the mixture was frozen and lyophilized to give the product (750 mg, 55%) as a red-brown solid. $^{1}$H NMR (500 MHz, DMSO-$d_6$) δ 1.04 (t, $J$ = 7.2 Hz, 3H), 1.43 (s, 3H), 1.44 (s, 3H), 2.31 (s, 3H), 2.50 (6H), 2.71 (s, 6H), 2.80 (s, 4H), 2.91 (t, $J$ = 6.0 Hz, 2H), 3.15 - 3.20 (m, 2H), 3.28 – 3.33 (s, 2H), 3.44 – 3.53 (m, 14H), 3.70 (t, $J$ = 5.9 Hz, 2H), 7.05 – 7.08 (m, 2H), 7.17 (d, $J$ = 8.0 Hz, 1H) 8.05 – 8.10 (m, 2H). $^{13}$C NMR (125 MHz, DMSO-$d_6$) δ 12.6, 12.7, 13.2, 14.7, 18.4, 25.4, 31.6, 33.6, 38.6, 43.6, 47.5, 65.2, 68.8, 69.4, 69.6, 69.7, 69.8, 118.6, 125.6, 126.6, 129.8, 130.0, 130.5, 132.1, 140.5, 140.7, 144.9, 153.7, 153.8, 153.9, 163.3, 163.6, 167.3, 170.1, 172.8.

### $^{1}H$-NMR Spectroscopy of ApHID and its precursors

PKS8324

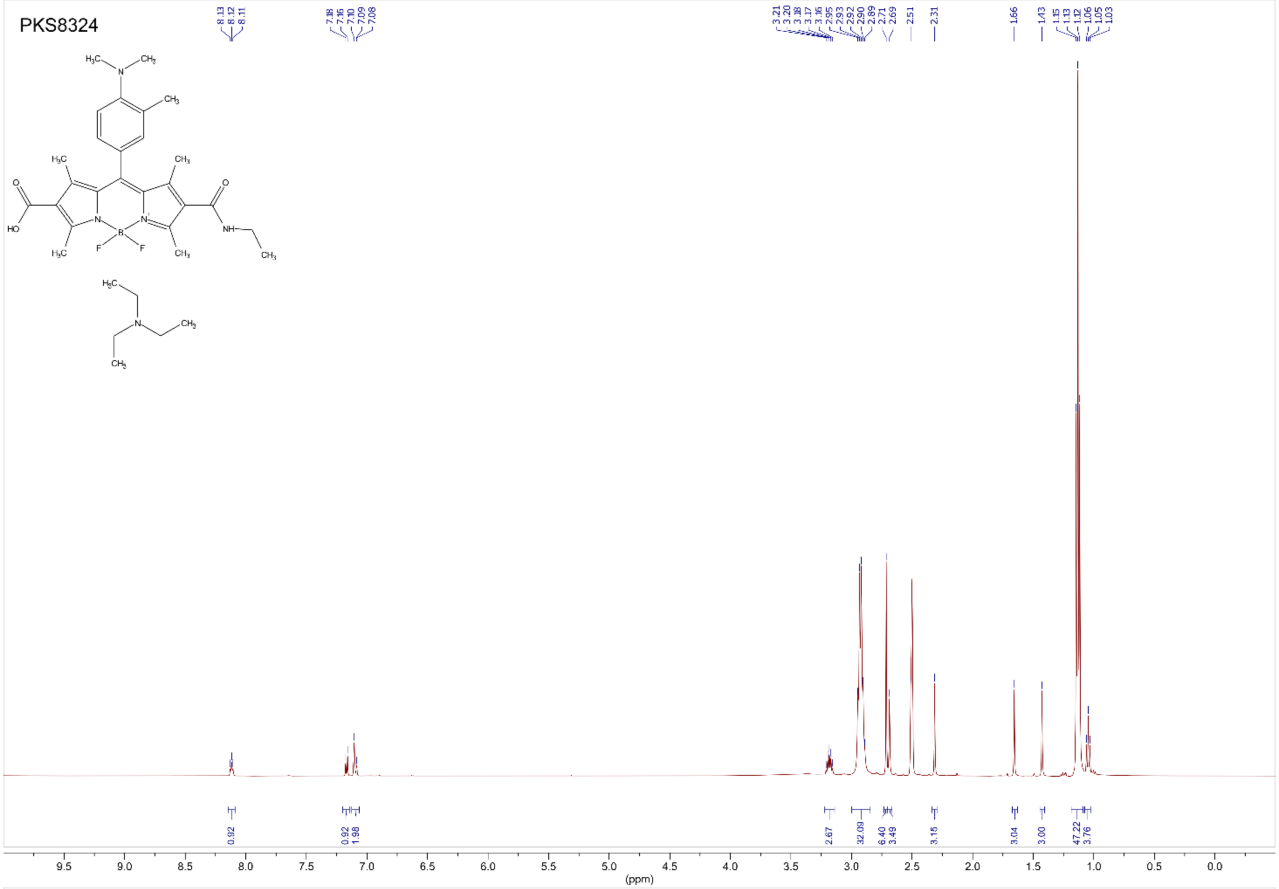

PKS8325

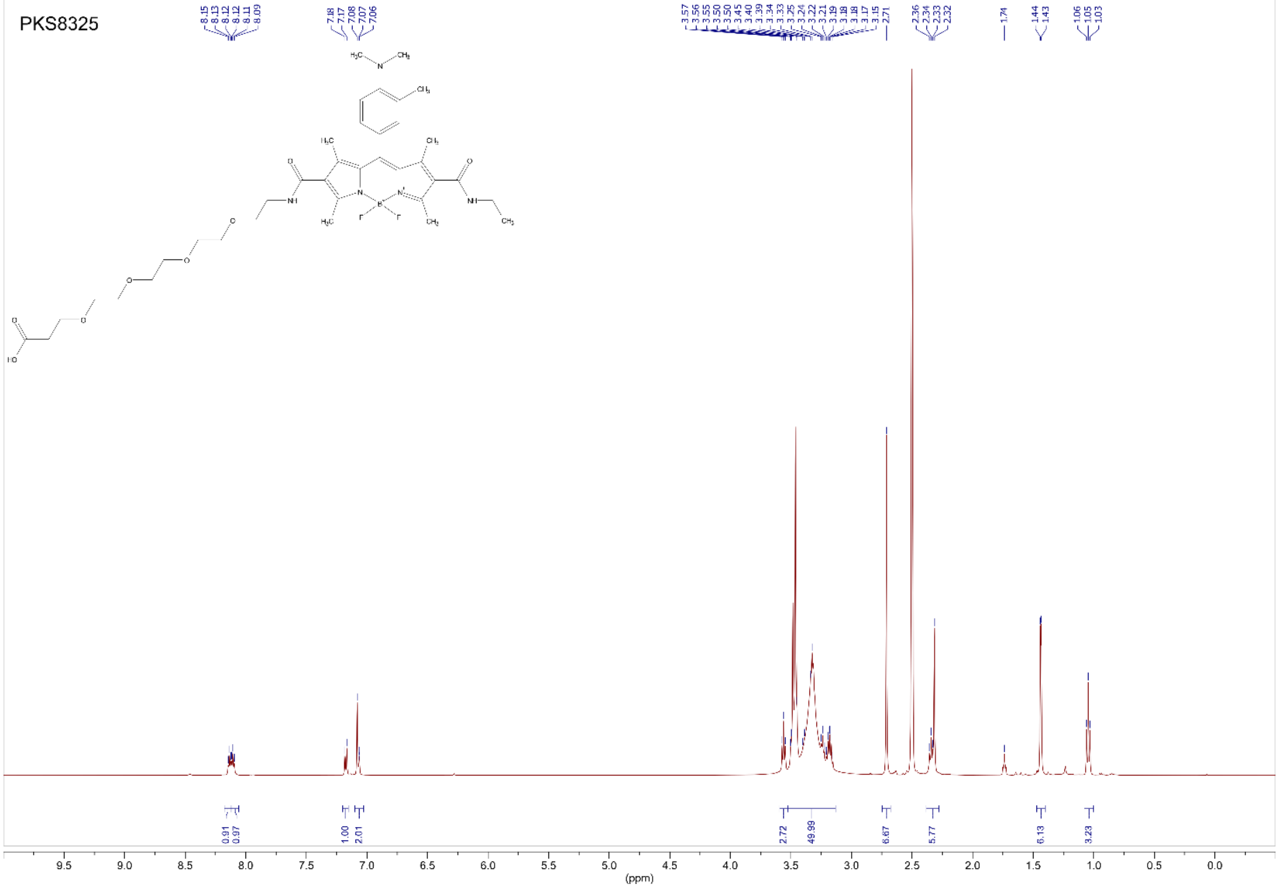

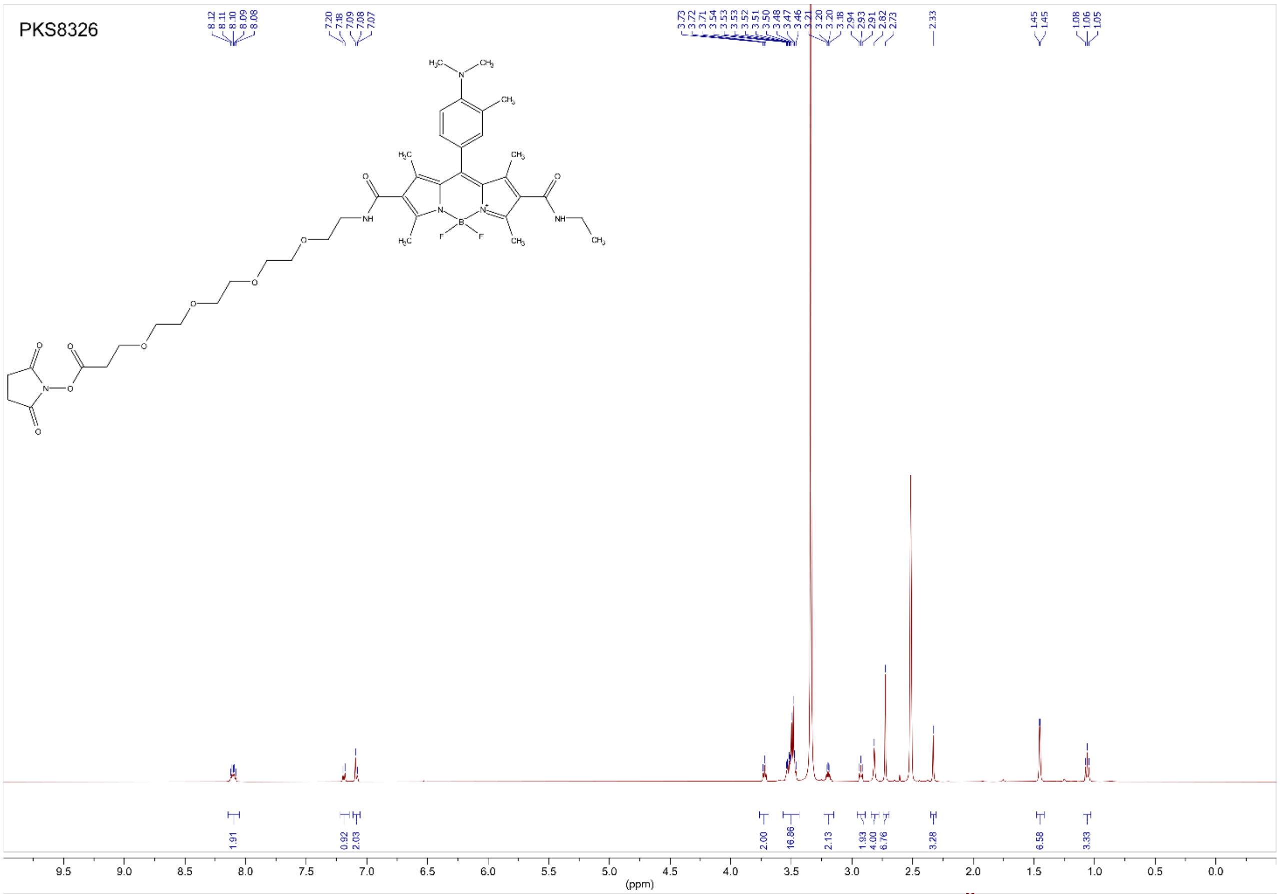
PKS8326
8.12
8.11
8.10
8.09
8.08
7.20
7.18
7.09
7.08
7.07
3.73
3.72
3.71
3.54
3.53
3.53
3.52
3.51
3.50
3.48
3.47
3.46
3.21
3.20
3.20
3.18
2.94
2.93
2.91
2.82
2.73
2.33
1.45
1.45
1.08
1.06
1.05
1.91
0.92
2.03
2.00
16.86
2.13
1.93
4.00
6.76
3.28
6.58
3.33
(ppm)

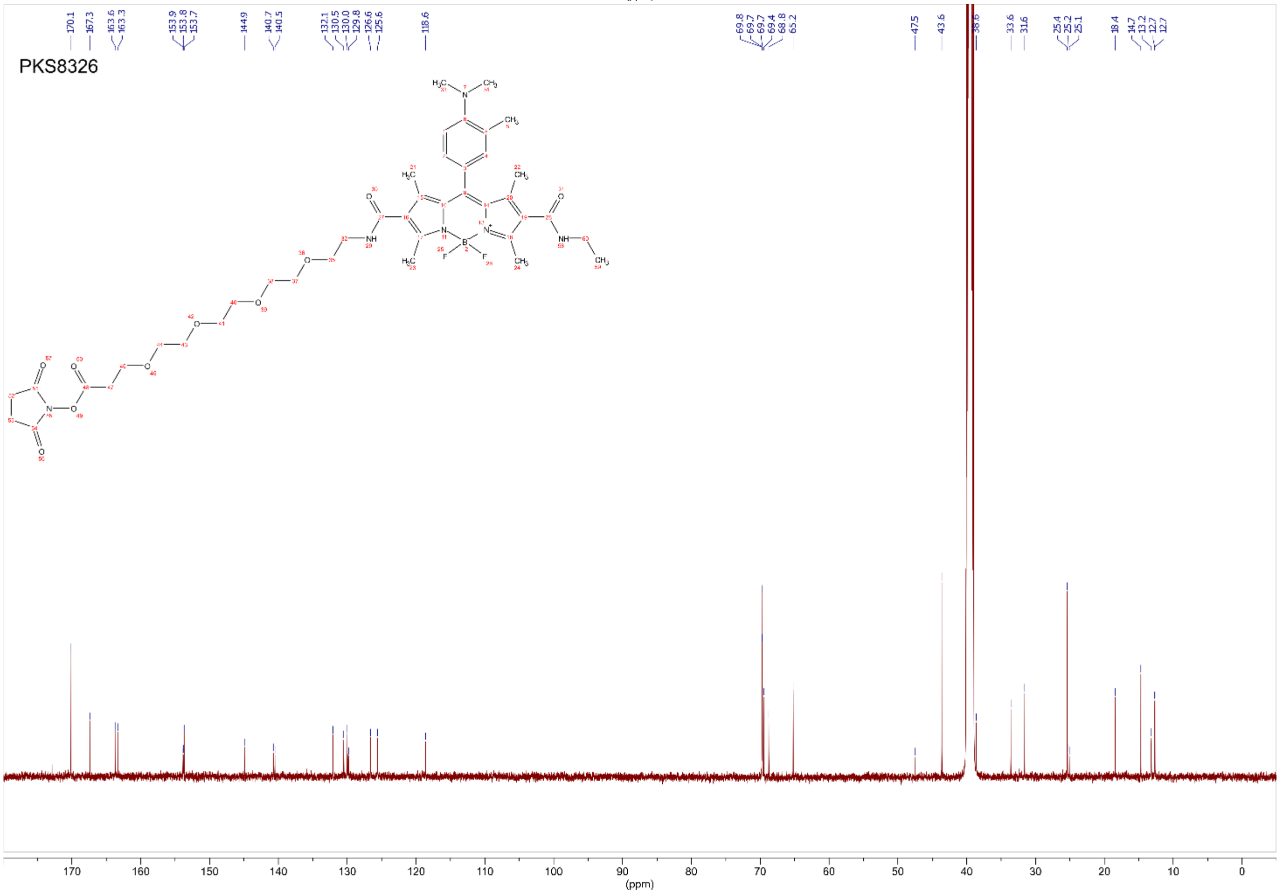
PKS8326
170.1
167.3
163.6
163.3
153.9
153.8
153.7
144.9
140.7
140.5
132.1
130.5
130.0
129.8
126.6
125.6
118.6
69.8
69.7
69.7
69.4
68.8
65.2
47.5
43.6
38.6
33.6
31.6
25.4
25.2
25.1
18.4
14.7
13.2
12.7
12.7
(ppm)

#### SUPPLEMENTARY REFERENCES


1. Maxfield, F.R. (1982). Weak bases and ionophores rapidly and reversibly raise the pH of endocytic vesicles in cultured mouse fibroblasts. J Cell Biol *95*, 676-681. 10.1083/jcb.95.2.676.

2. Nachliel, E., Finkelstein, Y., and Gutman, M. (1996). The mechanism of monensin-mediated cation exchange based on real time measurements. Biochim Biophys Acta *1285*, 131-145. 10.1016/s0005-2736(96)00149-6.

3. Gawinecki, R., Andrzejak, S., and Puchala, A. (1998). Efficiency of the Vilsmeier-Haack method in the synthesis of p-aminobenzaldehydes. Organic Preparations and Procedures International *30*, 455-460. Doi 10.1080/00304949809355310.

4. Li, M., Yao, Y., Ding, J., Liu, L., Qin, J., Zhao, Y., Hou, H., and Fan, Y. (2015). Spectroscopic and crystallographic investigations of novel BODIPY-derived metal-organic frameworks. Inorg Chem *54*, 1346-1353. 10.1021/ic502219y.

5. Maeda, H., Kowada, T., Kikuta, J., Furuya, M., Shirazaki, M., Mizukami, S., Ishii, M., and Kikuchi, K. (2016). Real-time intravital imaging of pH variation associated with osteoclast activity. Nat Chem Biol *12*, 579-585. 10.1038/nchembio.2096.
